# A high-resolution transcriptomic and spatial atlas of cell types in the whole mouse brain

**DOI:** 10.1101/2023.03.06.531121

**Authors:** Zizhen Yao, Cindy T. J. van Velthoven, Michael Kunst, Meng Zhang, Delissa McMillen, Changkyu Lee, Won Jung, Jeff Goldy, Aliya Abdelhak, Pamela Baker, Eliza Barkan, Darren Bertagnolli, Jazmin Campos, Daniel Carey, Tamara Casper, Anish Bhaswanth Chakka, Rushil Chakrabarty, Sakshi Chavan, Min Chen, Michael Clark, Jennie Close, Kirsten Crichton, Scott Daniel, Tim Dolbeare, Lauren Ellingwood, James Gee, Alexandra Glandon, Jessica Gloe, Joshua Gould, James Gray, Nathan Guilford, Junitta Guzman, Daniel Hirschstein, Windy Ho, Kelly Jin, Matthew Kroll, Kanan Lathia, Arielle Leon, Brian Long, Zoe Maltzer, Naomi Martin, Rachel McCue, Emma Meyerdierks, Thuc Nghi Nguyen, Trangthanh Pham, Christine Rimorin, Augustin Ruiz, Nadiya Shapovalova, Cliff Slaughterbeck, Josef Sulc, Michael Tieu, Amy Torkelson, Herman Tung, Nasmil Valera Cuevas, Katherine Wadhwani, Katelyn Ward, Boaz Levi, Colin Farrell, Carol L. Thompson, Shoaib Mufti, Chelsea M. Pagan, Lauren Kruse, Nick Dee, Susan M. Sunkin, Luke Esposito, Michael J. Hawrylycz, Jack Waters, Lydia Ng, Kimberly A. Smith, Bosiljka Tasic, Xiaowei Zhuang, Hongkui Zeng

**Author notes:** Corresponding authors: Zizhen Yao and Hongkui Zeng.

## Abstract

The mammalian brain is composed of millions to billions of cells that are organized into numerous cell types with specific spatial distribution patterns and structural and functional properties. An essential step towards understanding brain function is to obtain a parts list, i.e., a catalog of cell types, of the brain. Here, we report a comprehensive and high-resolution transcriptomic and spatial cell type atlas for the whole adult mouse brain. The cell type atlas was created based on the combination of two single-cell-level, whole-brain-scale datasets: a single-cell RNA-sequencing (scRNA-seq) dataset of ∼7 million cells profiled, and a spatially resolved transcriptomic dataset of ∼4.3 million cells using MERFISH. The atlas is hierarchically organized into five nested levels of classification: 7 divisions, 32 classes, 306 subclasses, 1,045 supertypes and 5,200 clusters. We systematically analyzed the neuronal, non-neuronal, and immature neuronal cell types across the brain and identified a high degree of correspondence between transcriptomic identity and spatial specificity for each cell type. The results reveal unique features of cell type organization in different brain regions, in particular, a dichotomy between the dorsal and ventral parts of the brain: the dorsal part contains relatively fewer yet highly divergent neuronal types, whereas the ventral part contains more numerous neuronal types that are more closely related to each other. We also systematically characterized cell-type specific expression of neurotransmitters, neuropeptides, and transcription factors. The study uncovered extraordinary diversity and heterogeneity in neurotransmitter and neuropeptide expression and co-expression patterns in different cell types across the brain, suggesting they mediate a myriad of modes of intercellular communications. Finally, we found that transcription factors are major determinants of cell type classification in the adult mouse brain and identified a combinatorial transcription factor code that defines cell types across all parts of the brain. The whole-mouse-brain transcriptomic and spatial cell type atlas establishes a benchmark reference atlas and a foundational resource for deep and integrative investigations of cell type and circuit function, development, and evolution of the mammalian brain.

## INTRODUCTION

The mammalian brain is arguably the most complex system in life, controlling a wide variety of organism’s activities including vitality, homeostasis, sleep, consciousness, sensation, innate behavior, goal-directed behavior, emotion, learning, memory, reasoning, and cognition. These activities are governed by highly specialized yet intricately integrated neural circuits in the brain. These circuits are composed of millions to billions of neurons and non-neuronal cells interconnected through a vast array of synaptic and non-synaptic intercellular communication machineries and molecules. These brain cells can be classified into numerous cell types based on various phenotypic measurements^1–5^. To understand how the variety of brain functions emerge from this complex system, it is essential to gain comprehensive knowledge about the cell types and circuits that constitute the molecular and anatomical architecture of the brain.

The anatomical architecture of the mammalian brain has been defined by its developmental plan and cross-species evolutionary ontology^6–8^. The entire brain is composed of telencephalon, diencephalon, mesencephalon (midbrain, MB), and rhombencephalon (hindbrain, HB).

Telencephalon consists of five major brain structures: isocortex, hippocampal formation (HPF), olfactory areas (OLF), cortical subplate (CTXsp) and cerebral nuclei (CNU). The first four brain structures, isocortex, HPF, OLF and CTXsp, constitute the developmentally derived pallium structure and are also collectively called cerebral cortex, whereas CNU derives from subpallium and is further divided into striatum (STR) and pallidum (PAL). Diencephalon consists of thalamus (TH) and hypothalamus (HY). Together telencephalon and diencephalon are also collectively referred to as forebrain. Hindbrain (HB) is divided into pons (P), medulla (MY), and cerebellum (CB). Within each of these major brain structures, there are multiple regions and subregions, each comprising many cell types.

Functionally, the mammalian brain is organized into four major systems: sensory, motor, cognitive and behavioral state^8^. Each of these systems contains multiple subsystems, which are organized in parallel and/or hierarchical manner across the above brain structures. The sensory system receives and processes sensory information from the periphery via multiple parallel ascending subsystems specific to different sensory modalities, i.e., visual, auditory, olfactory, taste, somatic, visceral, hormonal, and nociceptive. The motor system controls body function through the somatic, autonomic, and neuroendocrine subsystems. The motor system is generally organized in a hierarchical manner, with pools of motor neurons as the outputs that are controlled by several levels of central pattern generators, initiators, and controllers across the upstream regions of the brain. The cognitive system drives thinking and voluntary control of behaviors. It is also hierarchically organized, with cerebral cortex at the top followed by striatum and pallidum, all three levels interconnected via sequential descending projections. The cerebral cortex consists of multiple functionally specialized areas that form parallel circuit pathways with downstream regions. The behavioral state system comprises a series of localized cell groups, distributed in the ventral parts of the brain from cerebral nuclei to medulla, that control sleep and wakefulness and modulate behavioral states, often through the release of modulatory neurotransmitters and neuropeptides.

Cell types are considered the basic functional units of metazoan organs including the brain^4^, and they exhibit extraordinary diversity in their molecular, anatomical, physiological and functional properties. Significant progress has been made in characterizing these cellular properties and using them to classify cell types throughout the brain^1, 3–5, 9, 10^. Efforts have been dramatically accelerated by the advance of high-throughput single-cell genomics technologies over the past decade^4, 5^. Single-cell transcriptomics by single-cell or single-nucleus RNA sequencing (scRNA-seq or snRNA-seq) provides unprecedented depth of profiling and scalability, enabling comprehensive quantitative analysis and classification of cell types at scale^4, 5, 11–13^. This approach has been used to categorize cell types from many different regions of the mouse nervous system, such as cortex, hippocampus, striatum, thalamus, hypothalamus, cerebellum, spinal cord, and retina^14–29^, and increasingly more in human and non-human primate brains^30–38^. The BRAIN Initiative Cell Census Network (BICCN) and the Human Cell Atlas (HCA) are representative community efforts using single-cell transcriptomics to create cell type atlases for the brain and body of human and other mammals^12, 39–42^.

These studies have revealed important organizing principles of cell types in different parts of the brain, such as the hierarchical organization of cell types and the coexistence of discrete and continuous variation^4, 39^, as well as key gene networks related to cell type identities and structural/functional properties. In many cases, these studies have recapitulated previous sporadic knowledge about specific cell types, and further organized cell type information in an unmatched systematic and comprehensive manner. Furthermore, single-cell transcriptomic studies carried out in developing brains^43–52^ and in different species^30, 32, 38, 53–57^ have demonstrated that the transcriptomic cell type framework is a strong basis for elucidating the relationships between cell types that are rooted in their developmental and evolutionary origins.

An essential next step is to create a comprehensive and high-resolution transcriptomic cell type atlas for the entire adult brain from a single mammalian species. The mouse (*Mus musculus*) is the most widely used mammalian model organism and therefore a natural first choice for a comprehensive definition of mammalian brain composition and architecture. To define the anatomical context for cell types, another critical requirement is to obtain the precise spatial location of each cell type using single-cell-level spatial transcriptomics analysis^58–61^ covering the entire mouse brain. In addition to describing a complete, brain-wide cell type atlas of a mammalian brain, this analysis will provide essential knowledge about the cell type composition of different regions and circuits of the brain. The result is a foundational resource for conducting connectional and functional studies to understand how cell types interact to form neural circuits and what functional roles these cell types play, and for building additional cell type atlases across lifespan and for other species including human, to unravel the developmental and evolutionary bases of cell type organization and function.

As part of the BRAIN Initiative Cell Census Network (BICCN, www.biccn.org), we set out to build a comprehensive, high-resolution transcriptomic cell type atlas for the whole adult mouse brain, as a reference brain cell atlas for the neuroscience community. We generated a large-scale scRNA-seq dataset, with ∼7 million cells profiled across the entire mouse brain using the 10x Genomics Chromium platform, and several multiplexed error-robust fluorescence in situ hybridization (MERFISH)^62^ datasets covering the whole mouse brain. We conducted large-scale computational analysis of these datasets and derived a transcriptomic cell type taxonomy and atlas with ∼5,200 clusters organized into a hierarchical tree. The spatial locations of all the cell types were mapped in a cell atlas registered to the 3D Allen Mouse Brain Common Coordinate Framework version 3 (CCFv3)^63^ (**Supplementary Table 1** provides the anatomical ontology with full names and acronyms of all brain regions). We systematically characterized the distributions and relationships of all neuronal and non-neuronal cell types, identifying a high degree of correspondence between cell-type molecular profiles and their spatial distribution patterns. An investigation of transcription factor genes and related modules with specific expression at different hierarchical levels revealed their importance in defining cell types^64, 65^.

## RESULTS

### Creation of a high-resolution whole mouse brain transcriptomic cell type atlas

To create a high-resolution transcriptomic and spatial cell type atlas covering the entire mouse brain, we systematically generated two types of large-scale single-cell-resolution transcriptomic datasets for all mouse brain regions, by single-cell RNA-sequencing (scRNA-seq) and by MERFISH^62^, a spatially resolved transcriptomic method. We used the scRNA-seq datasets to generate a transcriptomic cell type taxonomy, and the MERFISH datasets to visualize and annotate the spatial location of each cluster in this taxonomy.

We first generated 781 scRNA-seq libaries (using 10x Genomics Chromium v2 or v3) from anatomically defined, CCFv3-guided (**Supplementary Table 1**) tissue microdissections (**Methods**), resulting in a dataset of ∼7.0 million single-cell transcriptomes (**Supplementary Table 2, 3**). We developed a set of stringent quality control (QC) metrics guided by pilot clustering results that informed us on characteristics of low-quality single-cell transcriptomes (**Methods**, **Supplementary Table 4**, **Extended Data Figure 1a-c**). We then conducted iterative clustering analysis on ∼4.3 million QC-qualified cells using custom software (scrattch.bigcat package developed in-house). The 10xv3 and 10xv2 cells were first clustered separately, and then integrated with methods we developed previously^28^, resulting in an initial joint transcriptomic cell type taxonomy with 5,283 clusters (**Extended Data Figure 1a**).

**Figure 1.**
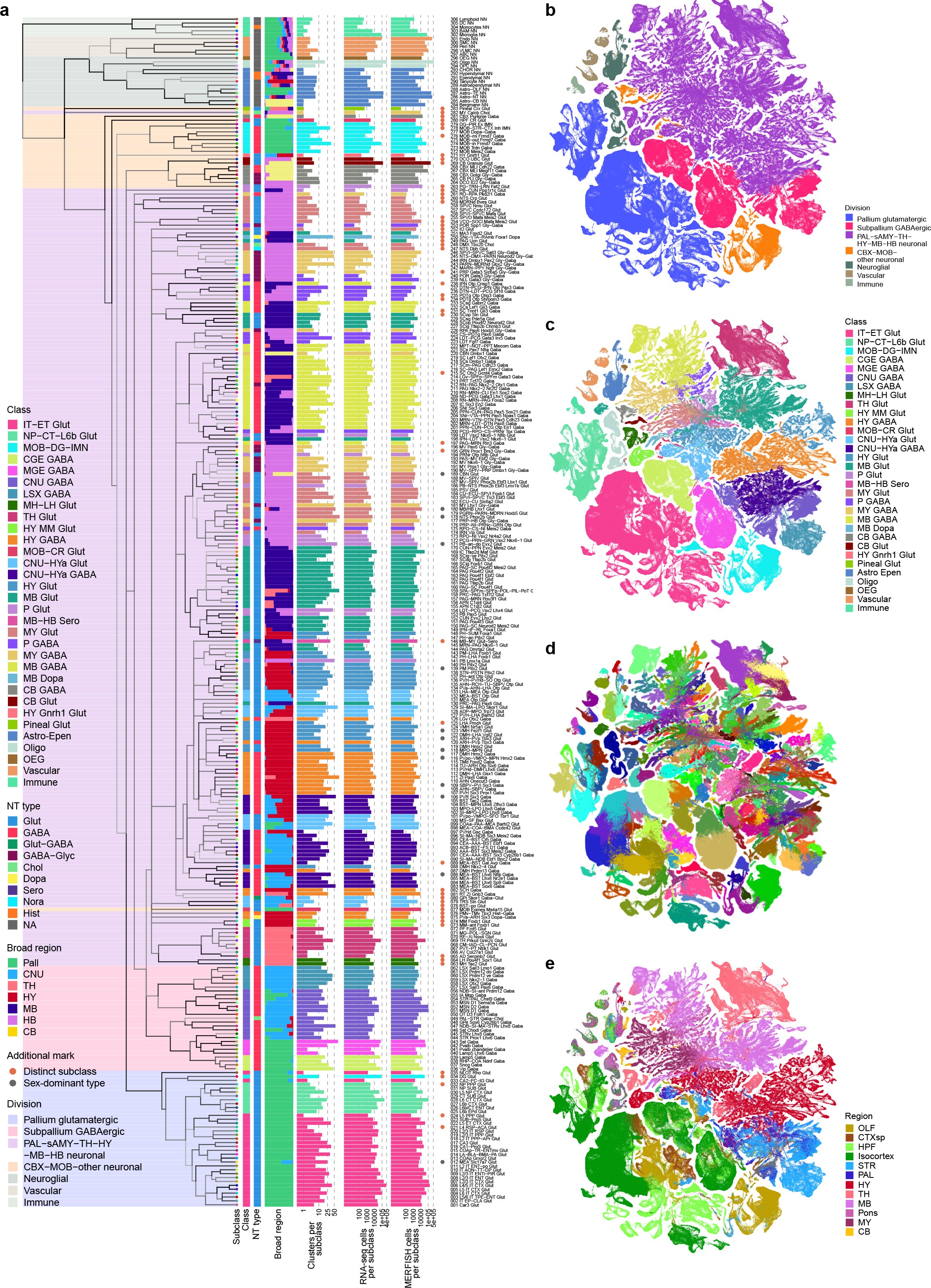
Transcriptomic cell type taxonomy of the whole mouse brain. (a) The transcriptomic taxonomy tree of 306 subclasses organized in a dendrogram (10xv2: n = 1,708,450 cells; 10v3 n = 2,349,599 cells). The color blocks divide the dendrogram into major cell divisions. From left to right, the bar plots represent class, major neurotransmitter type, region distribution of profiled cells, number of clusters, number of RNA-seq cells, and number of MERFISH cells per subclass. The subclasses marked with orange dots represent highly distinct subclasses and ones marked with grey dots represent subclasses containing sex-dominant clusters. For each cell, 15 nearest neighbors in reduced dimension space were determined and summarized by subclass. Highly distinct subclasses were identified as those with no nearest neighbors assigned to other subclasses and/or those that formed a highly distinct branch on the taxonomy dendrogram. Sex-dominant clusters within a subclass were identified by calculating the odds and log P value for Male and Female distribution per cluster. Clusters with odds < 0.2 and logPval < -10 were marked as sex-dominant. **(b-e)** UMAP representation of all cell types colored by division (b), class (c), subclass (d), and brain region (e).

By performing all pair-wise cluster comparisons in this initial transcriptomic taxonomy, we derived 8,108 differentially expressed genes (DEGs, **Supplementary Table 5**) differentiating all pairs of clusters. We then designed two sets of gene panels for the generation of MERFISH data, with each gene panel containing a selected set of marker genes with the greatest combinatorial power to discriminate among all clusters. The first gene panel contained 1,147 genes and was used by the Zhuang lab to generate MERFISH datasets from several male and female mouse brains (see companion manuscript Zhang et al. for details) using a custom imaging platform. The second gene panel contained 500 genes (**Supplementary Table 6**) and was used to generate a MERFISH dataset from one male mouse brain at the Allen Institute for Brain Science (AIBS) using the Vizgen MERSCOPE platform (**Extended Data Figure 2**). The AIBS MERFISH dataset contained 59 serial full coronal sections at 200-µm intervals spanning the entire mouse brain, with a total of ∼4.3 million segmented and QC-passed cells (**Extended Data Figure 2**), subsequently registered to the Allen CCFv3 (**Methods**).

**Figure 2.**
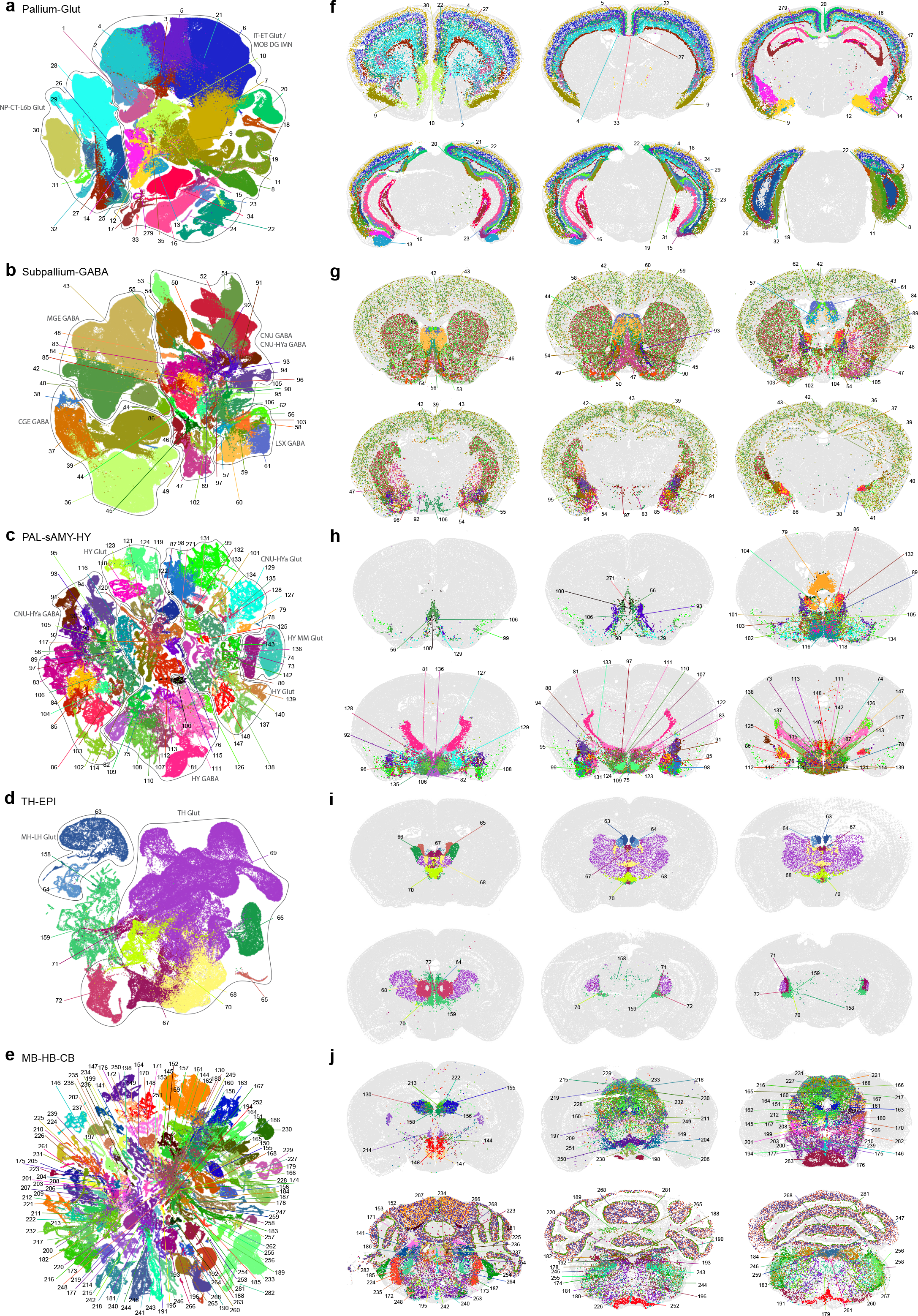
Neuronal cell type classification and distribution across the brain. UMAP representation **(a-e)** and representative MERFISH sections **(f-j)** of Pallium glut (a,f), Subpallium GABA (b,g), PAL-sAMY-HY (c,h), TH-EPI (d,i), and MB-HB-CB (e,j) neighborhoods colored by subclass. Each subclass is labeled by its ID and shown in the same color between UMAPs and MERFISH sections. Outlines in (a-d) show cell classes. For full subclass names see **Supplementary Table 7**.

To hierarchically organize the transcriptomic cell type taxonomy and better delineate the relationship between clusters, we computationally grouped the clusters into 306 subclasses (**Methods**). We used the AIBS MERFISH dataset and one of Zhuang lab’s MERFISH datasets to annotate the spatial location of each subclass and each cluster. To do this, we first mapped each MERFISH cell to the transcriptomic taxonomy and assigned the best matched cluster identity along with a correlation score to each MERFISH cell (**Methods**). The spatial location of each cluster was subsequently obtained by the collective locations of majority of the cells assigned to that cluster with high correlation scores. We annotated each subclass with its most representative anatomical region(s) and incorporated these annotations into subclass nomenclature for easier recognition of their identities. In this way, the high-level distribution of cell types across the entire mouse brain is described. As the anatomical annotations at subclass level are largely consistent between the Zhuang lab and the AIBS MERFISH datasets, in the subsequent sections of this manuscript, the AIBS MERFISH dataset is used to illustrate our results and findings.

To finalize the transcriptomic cell type taxonomy and atlas, we conducted detailed annotation and analysis of all the subclasses and clusters. During this process, we identified and removed an additional set of ‘noise’ clusters (usually doublets or mixed debris, see **Methods**) that had escaped the initial QC process, resulting in a final set of 5,200 high-quality clusters containing a total of ∼4.1 million high-quality single-cell transcriptomes (**Extended Data Figure 1a,d,e**).

Thorough analysis revealed extraordinarily complex relationships among transcriptomic clusters and their associated regions. Thus, to organize these complex molecular relationships, we derived a hierarchical representation of transcriptomic cell types (**Methods**). Overall, we defined a high-resolution transcriptomic and spatial cell type atlas for the whole mouse brain with 5 nested levels of classification: 7 divisions, 32 classes, 306 subclasses, 1,045 supertypes, and 5,200 clusters/types (**Table 1**, **Figure 1**, **Extended Data Figure 3**). **Supplementary Table 7** provides the cluster annotation, including the identity of the division, class, subclass and supertype assignment for each cluster, as well as full names of all levels of cell types and various metadata information. We provide several representations of this atlas for further analysis: a) a dendrogram at subclass resolution along with bar graphs displaying various metadata information (**Figure 1a**, **Extended Data Figure 3d**), b) UMAPs at single-cell resolution colored with different types of metadata information (**Figure 1b-e**, **Extended Data Figure 3c**), and c) a constellation diagram at subclass resolution to depict multi-dimensional relationships among different subclasses (**Extended Data Figure 4**).

**Figure 3.**
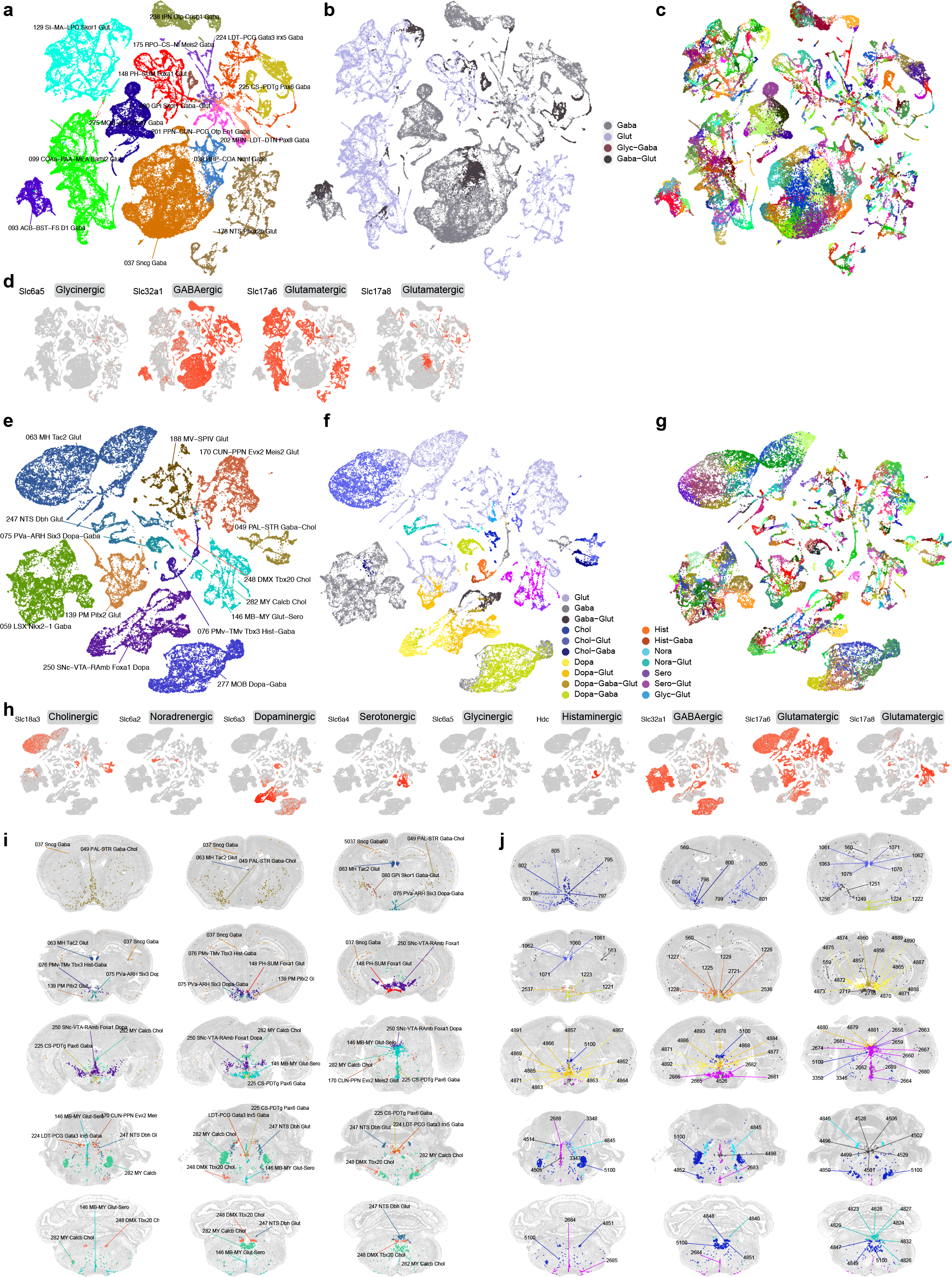
Neurotransmitter types and their distribution throughout the mouse brain. (a-c) UMAP representation of neuronal subclasses containing clusters releasing glutamate-GABA dual transmitters. UMAPs are colored by subclass (a), neurotransmitter type (b), and cluster (c). Glutamate-GABA co-releasing clusters include clusters *559*, *560*, *563* in subclass 37, cluster 566 in subclass 38, clusters *1249*, *1250*, *1251* in subclass 80, clusters 1498, 1499 in subclass 93, clusters 1571, 1592, 1593 in subclass 99, clusters 2307, 2308 in subclass129, clusters *2716*, *2717*, *2721* in subclass 148, clusters 3469, 3480, 3482 in subclass 175, cluster 3609 in subclass 178, cluster 4073 in subclass 201, cluster 4089 in subclass 202, clusters *4496*, *4498*, *4499*, *4501*, *4502*, *4505*, *4506*, *4514* in subclass 224, clusters *4526*, *4528*, *4529* in subclass 225, cluster 4653 in subclass 238, and cluster 5041 in subclass 275. Clusters in italic are shown in MERFISH sections in (j). **(d)** UMAPs representing the expression of neurotransmitter transporter genes for glutamate, GABA and glycine. **(e-g)** UMAP representation of neuronal subclasses containing clusters releasing modulatory neurotransmitters and their various combinations of co-releasing with glutamate and/or GABA. UMAPs are colored by subclass (e), neurotransmitter type (f), and cluster (g). Cholinergic neurons include clusters *795* (co-release w/ GABA), *796*, *797* (w/ GABA), *798* (w/ GABA), *799* (w/ glut), *800* (w/ GABA), *801*, *802*, *803-805* (all w/ glut) in subclass 49; cluster 958 (w/ GABA) in subclass 59; clusters *1060-1063*, *1070*, *1071* and *1075* (all w/ glut) in subclass 63; clusters 3322, *3346*, *3347*, *3348*, 3349, *3350* (all w/ glut except 3349) in subclass 170; cluster 3939 (w/ glut) in subclass 188; clusters *4847-4852* in subclass 248; and cluster *5100* in subclass 282. Dopaminergic neurons include clusters *1221-1224* (all w/ GABA) in subclass 75; clusters *2536* and *2537* (both w/ glut) in subclass 139; clusters *4856* (w/ glut- GABA), *4857* (w/ glut-GABA), *4860* (w/ glut-GABA), *4862* (w/ glut), *4863-4865*, *4866* (w/ glut), *4867*, *4868* (w/ glut), *4869* (w/ glut), *4870* (w/ GABA), *4871-4875*, *4876* (w/ GABA), *4877-4880* (all w/ glut), *4881*, *4883-4886* (all w/ glut), *4887-4890*, *4891* (w/ glut-GABA), *4892*, *4893* in subclass 250; and clusters 5047, 5048, 5050, 5055 (all w/ GABA) in subclass 277. Histaminergic neurons include clusters *1225* (w/ GABA), *1226* (w/ GABA), and *1227-1229* in subclass 76. Noradrenergic neurons include clusters *4823*, *4824*, *4826-4829*, *4832* and *4840* (all w/ glut), as well as *4845* in subclass 247. Serotonergic neurons include clusters *2658*, *2659*, *2660-2662* (all w/ glut), *2663*, *2664*, *2665-2667* (all w/ glut), *2674* (w/ glut), *2680*, *2681-2685* (all w/ glut), *2688* (w/ glut), and *2689* (w/ glut) in subclass 146. Clusters in italic are shown in MERFISH sections in (j). **(h)** UMAPs representing the expression of genes for glutamate, GABA and modulatory neurotransmitters. **(i-j)** Representative MERFISH sections showing the location of neuronal types with glutamate-GABA dual transmitters and those with modulatory neurotransmitters. Cells in (i) are colored and labeled by subclasses. Cells in (j) are colored by neurotransmitter/neuromodulator types and labeled by cluster IDs. See **Supplementary Table 7** for detailed neurotransmitter assignment for each cluster.

**Figure 4.**
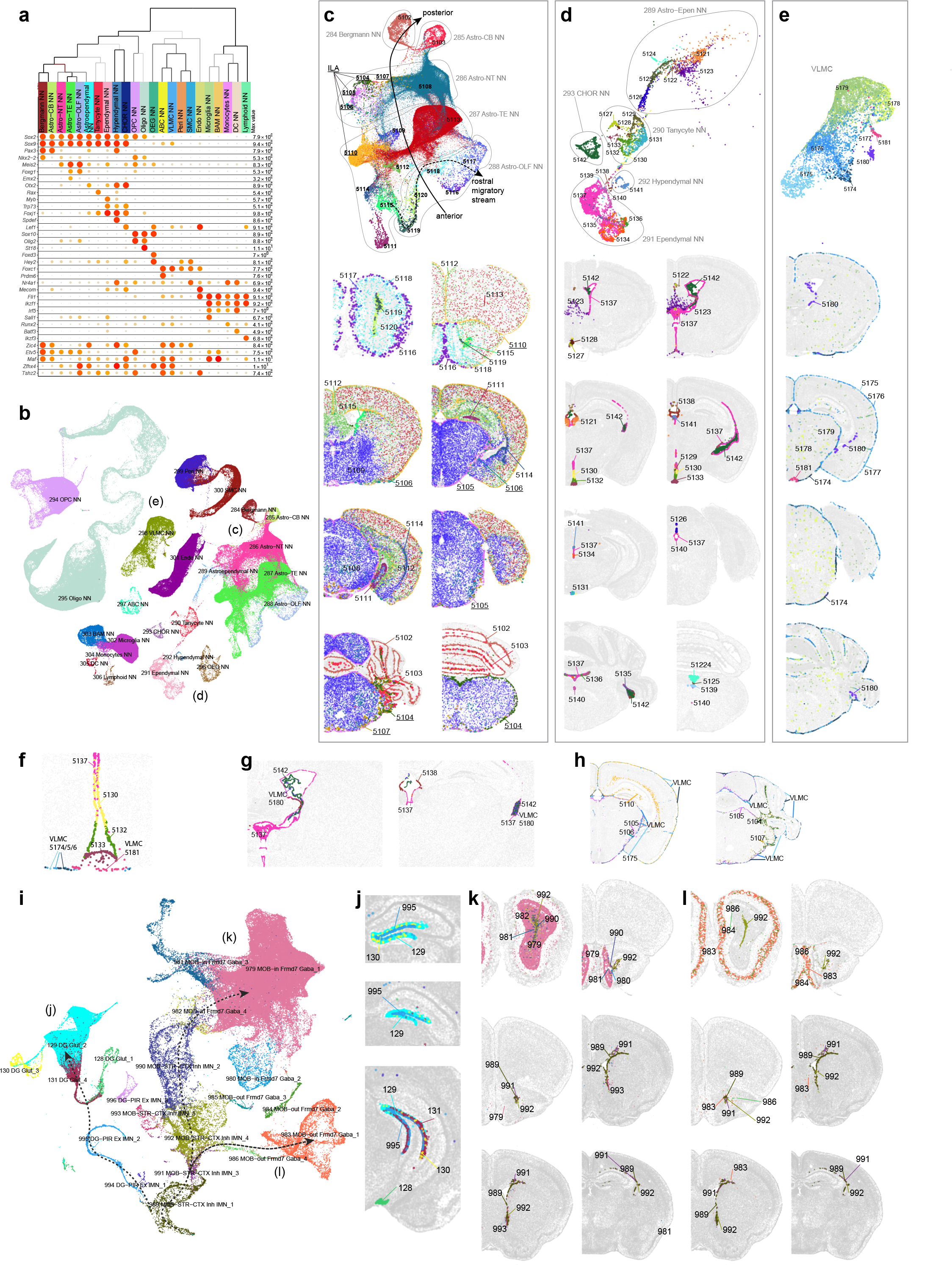
Non-neuronal cell types and immature neuronal types. (a) Dot plot showing the transcription factor marker gene expression in non-neuronal subclasses. Dot size and color indicate proportion of expressing cells and average expression level in each subclass, respectively. **(b)** UMAP representation of non-neuronal cell types colored by subclass. Three subpopulations are highlighted and further investigated: astrocytes (c), ependymal cells (d), and VLMC (e). **(c-e)** UMAP representation and representative MERFISH sections of astrocytes (c), ependymal cells (d), and VLMC (e) colored and numbered by cluster. Outlines in (c-d) UMAPs show subclasses. **(f)** Co-localization of VLMC cluster 5181 with Tanycyte cluster 5133 on the MERFISH section. **(g)** Co-localization of VLMC cluster 5180 with CHOR cluster 5142 and Ependymal clusters 5137 and 5138. **(h)** Co-localization of VLMCs with Interlaminar astrocytes (ILA). **(i)** UMAP representation of immature neuron populations colored by supertype. Maturation trajectories in dentate gyrus (DG) (j), inner main olfactory bulb (k), and outer main olfactory bulb (l) are highlighted. **(j-l)** Representative MERFISH sections showing location of immature neuronal supertypes from the three trajectories.

**Table 1.** Summary of the whole mouse brain cell type atlas. Major cell divisions (Pallium glutamatergic, Subpalium GABAergic, PAL-sAMY-TH-HY-BM-HB neuronal, CBX-MOB- other neuronal, Neuroglial, Vascular, Immune), cell classes under each division, and the numbers of subclasses, supertypes, and clusters under each class are listed. Each level of the hierarchy is color coded consistently with the taxonomy.

| Cell Class | No. of Subclasses | No. of Supertypes | No. of Clusters |
| --- | --- | --- | --- |
| **Pallium glutamatergic** |  |  |  |
| IT-ET Glut | 26 | 101 | 402 |
| NP-CT-L6b Glut | 8 | 27 | 83 |
| **Subpallium GABAergic** |  |  |  |
| CGE GABA | 4 | 19 | 101 |
| MGE GABA | 4 | 16 | 106 |
| CNU GABA | 13 | 50 | 212 |
| LSX GABA | 6 | 30 | 146 |
| **PAL-sAMY-TH-HY-MB-HB neuronal** |  |  |  |
| MH-LH Glut | 2 | 9 | 35 |
| TH Glut | 8 | 35 | 113 |
| CNU-HY<sub>a</sub> GABA | 18 | 83 | 379 |
| HY GABA | 18 | 72 | 425 |
| CNU-HY<sub>a</sub> Glut | 13 | 42 | 236 |
| HY Glut | 19 | 84 | 337 |
| HY MM Glut | 2 | 3 | 13 |
| MB Glut | 30 | 102 | 657 |
| P Glut | 11 | 38 | 235 |
| MY Glut | 22 | 59 | 411 |
| MB GABA | 26 | 87 | 472 |
| P GABA | 12 | 23 | 141 |
| MY GABA | 18 | 54 | 347 |
| MB Dopa | 1 | 8 | 43 |
| MB-HB Sero | 1 | 7 | 32 |
| **CBX-MOB-other neuronal** |  |  |  |
| CB GABA | 6 | 12 | 27 |
| CB Glut | 2 | 3 | 9 |
| HY Gnrh1 Glut | 1 | 1 | 1 |
| MOB-DG-IMN | 9 | 28 | 121 |
| MOB-CR Glut | 2 | 6 | 16 |
| Pineal Glut | 1 | 1 | 1 |
| **Neuroglial** |  |  |  |
| Astro-Epen | 10 | 22 | 41 |
| Oligo | 2 | 6 | 24 |
| OEG | 1 | 1 | 4 |
| **Vascular** |  |  |  |
| Vascular | 5 | 8 | 19 |
| **Immune** |  |  |  |
| Immune | 5 | 8 | 11 |

The high quality of the scRNA-seq data included in the final taxonomy is indicated by the high gene and UMI counts across the cell divisions (**Extended Data Figure 3a,b**). To test the robustness of the clustering results, we first performed 5-fold cross-validation using all 8,108 markers as features for classification to assess how well the cells could be mapped to the cell types they were originally assigned to. The median classification accuracy is 0.86 ± 0.10 (median ± SD) and 0.97 ± 0.03 for all clusters and all subclasses respectively. Next, we evaluated the integration between 10xv2 and 10xv3 transcriptomes. The UMAP shows good inter-mixing of 10xv2 and 10xv3 transcriptomes overall (**Extended Data Figure 5a-c)**. For cell types/clusters containing many cells, we observed separation of 10xv2 and 10xv3 data in the UMAP space, but not at the cluster level. For each of the 5,383 marker genes shared between 10xv2 and 10xv3 datasets, we computed the Pearson correlation of its average expression in each cluster for all overlapping clusters between the 10xv2 and 10xv3 data (**Extended Data Figure 5d**). The median correlation is 0.89 ± 0.09, suggesting a majority of the marker genes show consistent relative expression levels across clusters between the two 10x platforms. We manually inspected several genes with poor correlation and found them to have poor gene annotation or show relatively small variations across clusters. Lastly, we examined consistency of gene expression between 10xv3 and MERFISH data in corresponding cell types in a similar way and found high median Pearson correlation at 0.91 ± 0.15 (**Extended Data Figure 5d**). Most genes with low correlations are *Rik genes that are more likely to be poorly annotated, and the MERFISH probes selected for them might not work well. The MERFISH dataset can resolve the vast majority of clusters due to strong correlation of DEG expression between 10xv3 and MERFISH clusters (**Extended Data Figure 5e-g**). On the other hand, a few hundred pairs of clusters with fewer than two DEGs on the MERFISH gene panel remain unresolvable in the MERFISH data, and they are usually sibling clusters with indistinguishable spatial distribution.

**Figure 5.**
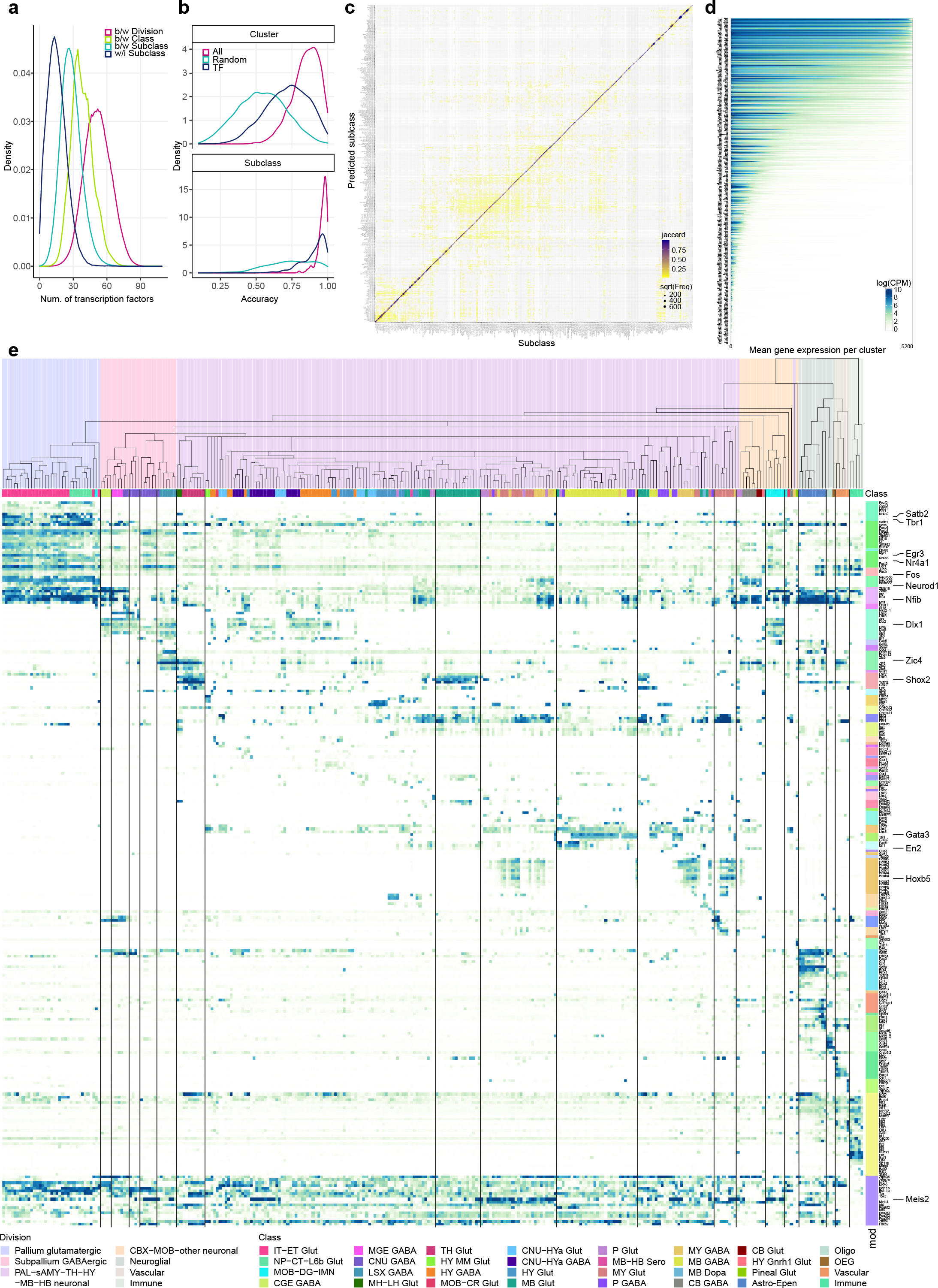
Transcription factor modules across the whole mouse brain. (a) Distribution of the number of differentially expressed TFs between divisions (pink), between classes (apple green), between subclasses (sea green), and within subclasses (dark blue). **(b)** Cross validation accuracy for each cluster (top panel) or subclass (bottom panel) using classifiers built based on all 8,108 marker genes (pink), randomly selected 499 marker genes (sea green), or 499 TF marker genes (dark blue). **(c)** Confusion matrix between the assigned and predicted subclasses using classifiers trained on 499 TF markers in cross validation. The size of the dots corresponds to the number of overlapping cells, and the color corresponds to the Jaccard similarity score between the assigned and predicted subclasses. **(d)** Expression level of TFs (logCPM) per cluster. For each TF along the Y axis, clusters are sorted from the highest to lowest mean gene expression level along the X axis. **(e)** Expression of key TFs for each subclass in the taxonomy tree, organized in gene modules (mod) shown as color bars on the right. The color blocks divide the dendrogram into major cell divisions. The color bars below the dendrogram denote classes.

### Organization of neuronal cell types across the mouse brain

Neuronal cell types constitute a large proportion of the whole brain cell type atlas, including 4 divisions, 27 classes (84%), 283 subclasses (92%), 1,000 supertypes (95%) and 5,101 clusters (98%; **Table 1**, **Supplementary Table 7**). Neuronal types are distributed across all major brain structures, have high regional specificity, and exhibit highly variable degrees of similarities and differences amongst each other. Of the 4 neuronal divisions, glutamatergic neurons from all pallium structures, including isocortex, hippocampal formation (HPF), olfactory areas (OLF) and cortical subplate (CTXsp), form a distinct “Pallium glutamatergic” division (**Table 1**, **Figure 1a**, **Extended Data Figure 4**). Similarly, a set of developmental subpallium-derived GABAergic neuronal subclasses, including all GABAergic neurons found in pallium structures and those in the subpallial cerebral nuclei (CNU), including dorsal and ventral striatum (STRd and STRv), lateral septal complex (LSX), and dorsal, ventral and medial pallidum (PALd, PALv and PALm), form a second “Subpallium GABAergic” division (**Table 1**, **Figure 1a**, **Extended Data Figure 4**). We also identified a variety of distinct neuronal subclasses, including those from the main olfactory bulb (MOB) and cerebellar cortex (CBX), and tentatively grouped them into a mixed “CBX-MOB-other neuronal” division (**Table 1**, **Figure 1a**, **Extended Data Figure 4**).

Interestingly, in contrast to these highly distinct neuronal subclasses, the large set of remaining neuronal subclasses spanning the middle parts of the brain, including the striatum-like amygdala nuclei (sAMY) and pallidum (PAL) parts of CNU, thalamus (TH), hypothalamus (HY), midbrain (MB) and hindbrain (HB), exhibit a high degree of similarity and continuity, and hence were grouped into a single large “PAL-sAMY-TH-HY-MB-HB neuronal” division (**Table 1**, **Figure 1a**, **Extended Data Figure 4**).

To further investigate the neuronal diversity within each major brain structure, we generated re-embedded UMAPs for subsets of neuronal types within divisions and brain structures. The process of subdivision and UMAP re-embedding (in 2D and 3D) was iteratively applied at more detailed levels to reveal fine-grained relationships between neuronal types within and between brain regions. We name the various re-embedded groups of cell types ‘neighborhoods’ and use them for visualization and analysis purposes. The results shown in **Figure 2** reveal a striking correspondence between transcriptomic specificity and relatedness and spatial specificity and relatedness among the different neuronal subclasses.

In the Pallium glutamatergic division (subclasses 1-35, total 494 clusters), each neuronal subclass exhibits layer and/or region specificity (**Figure 2a,f**). We found that the homologous relationships of the different subclasses of glutamatergic neurons between isocortex and HPF we had reported previously^28^ extended to other pallium structures, i.e., OLF and CTXsp. We also observed that the NP-CT-L6b-like (NP: near-projecting, CT: corticothalamic, L6b: layer 6b) subclasses emerged as a group highly distinct from the IT-ET-like (IT: intratelencephalic, ET: extratelencephalic) subclasses^25, 27, 28, 39^. Thus, we defined two classes, IT-ET and NP-CT-L6b, for this division.

Based on the molecular signature and regional specificity of each subclass, the Subpallium GABAergic division (subclasses 36-62, total 565 clusters) was divided into four classes that are likely related to their distinct developmental origins^66, 67^ (**Figure 2b,g**): CGE GABA (containing cortical/pallial GABAergic neurons derived from the caudal ganglionic eminence), MGE GABA (containing cortical/pallial GABAergic neurons derived from the medial ganglionic eminence), CNU GABA (containing striatal/pallidal GABAergic neurons derived from the lateral ganglionic eminence, LGE, as well as from MGE and the embryonic preoptic area), and LSX GABA (containing lateral septum GABAergic neurons derived from the embryonic septum^68^).

We divided the large PAL-sAMY-TH-HY-MB-HB neuronal division (containing subclasses 63- 262 and 282, excluding 77, total 3873 clusters) into several neighborhoods to illustrate cell type organization in each major brain structure. The PAL-sAMY-HY neighborhood contains a set of closely related neuronal subclasses from the entire hypothalamus^24, 69^, as well as the sAMY and caudal PAL regions of CNU that are also known as the extended amygdala (**Figure 2c,h**). Both glutamatergic and GABAergic neuronal subclasses in this neighborhood exhibit a gradual anterior-to-posterior transition, and thus were grouped into five classes: CNU-HYa GABA, HY GABA, CNU-HYa Glut, HY Glut and HY MM Glut (MM standing for medial mammillary nucleus). Neuronal types in the most anterior part of HY, i.e., the preoptic area, are highly similar to neuronal types in sAMY and PAL. Some of the CNU-HYa GABA subclasses are also included in the Subpallium GABA neighborhood to show their relatedness and continuity with the striatal/pallidal types (**Figure 2b,g**). On the other hand, the more posterior HY GABA class also includes GABAergic neurons from the thalamic reticular nucleus (RT; subclass 81) and the ventral part of the lateral geniculate complex (LGv; subclass 126), which are closely related to zona incerta (ZI) neurons in HY (subclass 111), revealing a relationship of GABAergic types between hypothalamus and thalamus.

The TH-EPI neighborhood (**Figure 2d,i**) contains all glutamatergic neuronal subclasses from the thalamus, as well as the medial and lateral habenula (MH and LH) which collectively compose the epithalamus (EPI). These subclasses were grouped correspondingly into TH Glut and MH-LH Glut classes, except for one subclass with neurons found in several posterior thalamic nuclei, 159_SPA-SPFm-SPFp-POL-PIL-PoT Glut, which belongs to the MB Glut class, revealing a relationship of glutamatergic types between thalamus and midbrain.

Finally, we show an example large neighborhood (**Figure 2e,j**) containing all the glutamatergic and GABAergic neuronal subclasses from MB and HB, which contains pons (P), medulla (MY) and cerebellum (CB; thus also including the CBX subclasses from the CBX-MOB-other neuronal division). In this highly complex neighborhood, we defined the following 10 classes based on transcriptomic relatedness and regional specificity: MB Glut, P Glut, MY Glut, MB GABA, P GABA, MY GABA, MB dopa, MB-HB Sero, CB GABA and CB Glut (**Supplementary Table 7**). We found that the glutamatergic and GABAergic subclasses, 189 and 220, from the cerebellar nuclei (CBN) are more closely related to those from the medulla than those from the cerebellar cortex.

The analysis presented thus far provides a high-level overview of the extraordinary complexity of neuronal cell types across the brain. These data and the whole brain atlas will allow for more in-depth analyses to understand the relationship of neuronal types in different brain structures. Here, we also highlight a small set of remarkable neuronal types (defined at subclass level) that are transcriptomically highly distinct from all the other subclasses (**Extended Data Figure 6**, also marked with orange dots in **Figure 1a** and with red circles in **Extended Data Figure 4**).

**Figure 6.**
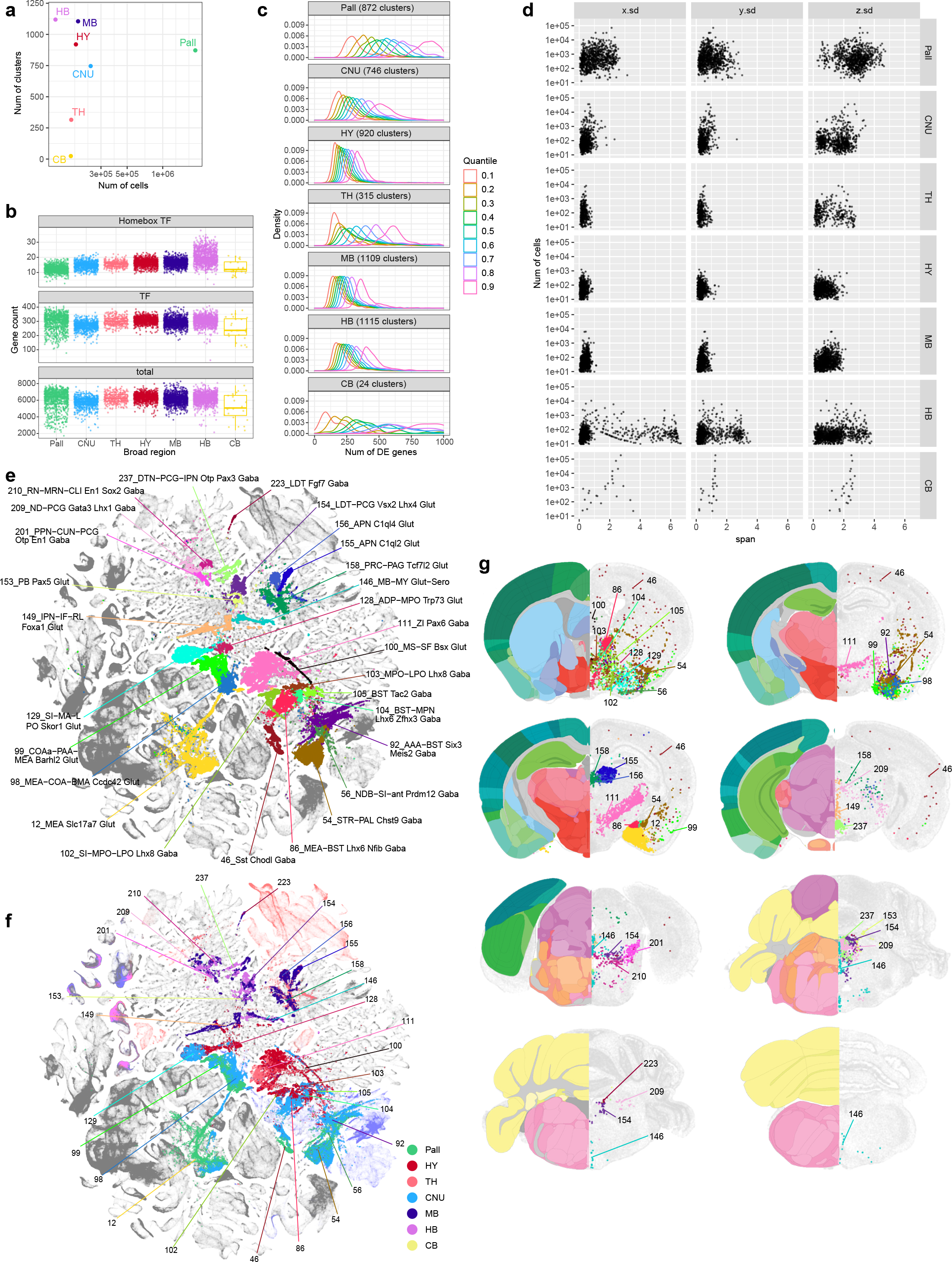
Region specific features and transitional cell types. (a) Scatterplot showing the number of neuronal clusters identified per region vs. the number of neuronal cells profiled within the corresponding region. Each neuronal cluster is assigned to the most dominant region. **(b)** Distribution of the number of genes detected per neuronal cluster per region with logCPM > 3. The top panel shows the number of Homeobox TFs per cluster per region, the middle panel shows the number of all TFs expressed per cluster per region, and the bottom panel shows the number of any gene expressed per cluster per region. **(c)** Distribution of the number of DEGs between every pair of neuronal clusters within each region, split at quantiles of 0.1, 0.2, …, and 0.9. The curves show the spread of the number of DEGs between more similar types at 0.1 quantile vs. the more distinct types at 0.9 quantile. **(d)** Scatterplot showing the number of cells mapped to a given neuronal cluster vs. the standard deviation of their 3D coordinates along the X (medial-lateral), Y(dorsal-ventral), and Z (anterior-posterior) axis based on the MERFISH dataset, stratified by the regions. The plot shows how localized the clusters are within each region along each spatial axis. **(e-g)** UMAP representation (e-f) and representative MERFISH sections (g) of subclasses shared between broad regions, (e,g) colored by subclass, and (f) colored by region. In (g) the best matching CCF reference atlas is shown on the left side of the MERFISH sections.

These highly distinct neuronal subclasses are found in all parts of the brain, each at a very specific anatomical location (**Extended Data Figure 6**). For example, subclass 21 is a L4 neuronal type with mixed IT and ET transcriptomic signatures in the retrosplenial cortex (RSP). Subclass 35 is a highly distinct IT-ET type located in the nucleus of the lateral olfactory tract (NLOT). Subclass 79 is located in the triangular nucleus of septum (TRS) specifically. Subclass 125 is located in the lateral hypothalamic area (LHA) and specifically expresses the neuropeptide gene *Pmch*. Subclass 230 is a superior colliculus (SC) glutamatergic type highly distinct from all the other SC neuronal types. Subclasses 234 and 235 are located in the posterodorsal tegmental nucleus (PDTg) specifically. Subclass 238 is primarily located in interpeduncular nucleus (IPN). Subclass 252 is specific to inferior olivary complex (IO). Subclass 263 is predominantly located in pontine gray (PG). Subclass 271 is the hypothalamic Gnrh1 neuronal type developmentally originated from the embryonic olfactory epithelium^70^. Subclass 281 is the cerebellar Purkinje neurons.

### Neurotransmitter identities and neuropeptide expression patterns in neuronal cell types

We systematically assigned neurotransmitter identity to each cell cluster based on the expression of canonical neurotransmitter transporter genes (**Figure 3**, **Extended Data Figure 3c-d**, **Supplementary Table 7**), i.e., *Slc17a7* (also known as *Vglut1*), *Slc17a6* (*Vglut2*) and *Slc17a8* (*Vglut3*) for glutamatergic, *Slc32a1* (*Vgat*) for GABAergic, *Slc6a5* for glycinergic, *Slc18a3* (*Vacht*) for cholinergic, *Slc6a3* (*Dat*) for dopaminergic, *Slc6a4* (*Sert*) for serotonergic, and *Slc6a2* (*Net*) for noradrenergic. The only exception was the use of the *Hdc* gene to identify histaminergic cells since there is no known high-affinity reuptake system for histamine^71^. We used a stringent expression threshold of log(CPM) > 3.5 of these genes to assign neurotransmitter identity to each cluster. We also used two commonly used marker genes, *Chat* for cholinergic neurons and *Dbh* for noradrenergic neurons, to further qualify or disqualify the assignments. For example, we found a few clusters that are *Slc6a2*-positive but *Dbh*-negative, and thus did not assign them the noradrenergic identity.

Based on these marker genes, the majority of neuronal clusters express a single neurotransmitter, either glutamate or GABA. Many GABAergic neuronal clusters in MB and HB co-express glycine. We identified 49 clusters with glutamate-GABA dual-transmitters (Glut-GABA), most of which utilize *Slc17a6* or *Slc17a8* as the glutamate transporter (**Supplementary Table 7**, **Figure 3a-d,i,j**). These clusters are widely distributed in different parts of the brain. They include 4 clusters in the isocortex and hippocampus and 3 clusters in globus pallidus, internal segment (GPi), which likely correspond to previously well-characterized glutamate-GABA co-releasing neuronal types in these regions^72, 73^. They also include a few clusters each in the cortical amygdala areas, STRv, ventral PAL, posterior HY, several MB areas including the ventral tegmental area (VTA), pedunculopontine nucleus (PPN) and interpeduncular nucleus (IPN), areas in pons such as superior central nucleus raphe (CS), nucleus raphe pontis (RPO) and laterodorsal tegmental nucleus (LDT), etc. (**Figure 3a-d,i,j**). Interestingly, except for the 3 glutamate-GABA clusters that form an exclusive subclass in GPi, the other Glut-GABA clusters are present in subclasses that also contain closely related single-neurotransmitter (glutamate or GABA) clusters (**Figure 3a-d**, **Supplementary Table 7**), and our QC process determined that this was not due to data quality issues (doublets or low-quality cells).

We also systematically identified all clusters producing modulatory neurotransmitters (**Figure 3e-j**, **Supplementary Table 7**). Cholinergic neurons^74, 75^ are found mainly in subclass 49 in the ventral PAL (10 clusters), but also include 1 cluster in LSX, 7 clusters in MH, 6 clusters in PPN and cuneiform nucleus (CUN), 6 clusters in dorsal motor nucleus of the vagus nerve (DMX), and 3 clusters scattered in other subclasses in MY. We also found *Slc18a3* expression in several clusters in the Vip GABA subclass, but its expression at cluster level did not cross our threshold to label these clusters as cholinergic. Cholinergic neurons often co-express glutamate (18 out of 34 clusters) or sometimes GABA (5 out of 34).

Dopaminergic neurons^18^ are found predominantly in subclass 250, which is the sole member of the MB Dopa class, located in substantia nigra, compact part (SNc), VTA and midbrain raphe nuclei (RAmb) areas. This subclass displays the most heterogeneous neurotransmitter content, consistent with previous findings^76^. It contains 34 dopaminergic clusters, as well as 9 glutamatergic, GABAergic or dual glutamate-GABA clusters. Some (18) of the 34 dopaminergic clusters also co-express glutamate, or GABA, or both glutamate and GABA. Dopaminergic neurons are also found in 6 clusters in arcuate hypothalamic nucleus (ARH) and ventral premammillary nucleus (PMv) of HY (co-expressed with GABA or glutamate) and in 4 clusters in MOB (co-expressed with GABA).

Serotonergic neurons^19^ all belong to the distinct MB-HB Sero class, which contains a single subclass, 146. This subclass consists of 19 serotonergic clusters and 13 glutamatergic (*Slc17a8*) clusters that are all closely related to each other. Some (14) of the 19 serotonergic clusters also co-express glutamate (*Slc17a8*). All these clusters reside in the various raphe nuclei within MB or MY. Thus, the serotonergic neuron class/subclass is highly heterogeneous in both neurotransmitter content and spatial localization.

Noradrenergic neurons^77, 78^ are mainly found in subclass 247. This subclass contains 9 noradrenergic clusters and 16 glutamatergic clusters, with all but one of the noradrenergic clusters also co-expressing glutamate (*Slc17a6*). All but two clusters in this subclass are located in the nucleus of the solitary tract (NTS), whereas the two exceptions (one noradrenergic and one glutamatergic cluster) are located in locus ceruleus (LC). Histaminergic neurons are found exclusively in the tuberomammillary nucleus, dorsal and ventral parts (TMd and TMv) of HY (5 clusters), two of which co-express GABA^71^. We found that ependymal and hypendymal cells, as well as monocytes may also be histaminergic (**Extended Data Figure 3d**).

Overall, an intriguing pattern emerged where nearly all subclasses with a dominant modulatory neurotransmitter contain clusters expressing glutamate and/or GABA only, as well as various forms of co-expression, indicating a high degree of heterogeneity in neurotransmitter release and co-release among closely related neuronal types that may have common developmental origins. Again, our QC process excluded the possibility of doublet or low-quality cell contamination.

While many of these transmitter co-release patterns had been documented previously^76, 79, 80^, our study defined a comprehensive set of cell types with unique and differing neurotransmitter content that can be tracked with marker genes.

Neuropeptides are also major agents for intercellular communications in the brain^81, 82^. We examined cell type-specific expression patterns of dozens of main neuropeptide genes and their receptors in our datasets (**Supplementary Table 7**). We measured the cell type specificity of expression of these genes using the Tau score^83^ and found a wide range of variation (**Extended Data Figure 7a-b**). Some neuropeptides are widely expressed in many cell types/clusters and at high levels (e.g., *Cck*, *Adcyap1*, *Pnoc*, *Penk*, *Sst* and *Tac1*), some are expressed at high levels in a moderate number of clusters (e.g., *Cartpt*, *Nts*, *Pdyn*, *Gal*, *Tac2*, *Grp*, *Vip*, *Crh*, *Trh* and *Cort*), whereas others are highly expressed specifically in only one or few clusters (e.g., *Avp*, *Agrp*, *Pomc*, *Pmch*, *Oxt*, *Rln3*, *Npw*, *Nps*, *Ucn*, *Hcrt*, *Gnrh1*, *Gcg* and *Pyy*; **Extended Data Figure 7c-f**). More than 80% of all clusters express at least one neuropeptide gene, and there are numerous co-expression combinations of different neuropeptides in many clusters, with high degrees of variations within subclasses (**Supplementary Table 7**). Our datasets provide a rich resource for the exploration of neuropeptide ligand/receptor interactions across the entire brain. However, we also note that the relationships between mRNA levels, the post-translationally processed peptide levels, and the functional levels are unknown for most neuropeptides, thus, it is difficult to predict what mRNA levels would lead to sufficient functional expression of a given neuropeptide (**Extended Data Figure 7c-d**).

### Non-neuronal cell types and immature neuron types across the mouse brain

The whole-brain transcriptomic cell type atlas describes the taxonomy of non-neuronal cell types, classifying them into 3 divisions (Neuroglial, Vascular and Immune), 5 classes, 23 subclasses, 45 supertypes and 99 clusters (**Table 1**, **Supplementary Table 7**), which can be distinguished by highly specific marker genes at all levels of hierarchy (**Figure 1a**, **Figure 4a-b**, **Extended Data Figure 8a-f**). The Neuroglial division comprises three classes, Astro-Epen, Oligo and OEG. The Astro-Epen class is the most complex, containing ten subclasses, four of which represent astrocytes that are specific to different brain regions: Astro-OLF, Astro-TE (for telencephalon), Astro-NT (for non-telencephalon) and Astro-CB, while the other six subclasses are astrocyte-related cell types: astroependymal cells, ependymal cells, tanycytes, hypendymal cells, choroid plexus (CHOR) cells, and Bergmann glia (**Figure 4a-d**). The Oligo class contains two subclasses, oligodendrocyte precursor cells (OPC) and oligodendrocytes (Oligo). The Oligo subclass is further divided into four supertypes corresponding to different stages of oligodendrocyte maturation: committed oligodendrocyte precursors (COP), newly formed oligodendrocytes (NFOL), myelin-forming oligodendrocytes (MFOL), and mature oligodendrocytes (MOL) (**Extended Data Figure 8h**). The OEG class corresponds to olfactory ensheathing glia (OEG). The Vascular division (and class) consists of five subclasses: arachnoid barrier cells (ABC), vascular leptomeningeal cells (VLMC), pericytes (Peri), smooth muscle cells (SMC), and endothelial cells (Endo). The Immune division (and class) is composed of five subclasses: microglia, border-associated macrophages (BAM), monocytes, dendritic cells (DC), and lymphoid cells, which contains B cells, T cells, NK cells and innate lymphoid cells (ILC).

We identified transcription factors (TFs) that potentially serve as master regulators for many of these non-neuronal cell types (**Figure 4a**, **Extended Data Figure 8d**), many of which were well documented in the literature^29, 84–88^. For example, *Sox2*, a well-known radial glia marker, is widely expressed in neuroglia, *Sox9* is specific to the Astro-Epen class, *Sox10* is specific to the Oligo class, *Foxd3* and *Hey2* are specific to OEG, *Foxc1* is specific to the Vascular division, and *Ikzf1* is specific to the Immune division. Within each division and class, additional TFs mark finer groupings^29^ (**Extended Data Figure 8d-f**). For example, Astro-TE cells express *Foxg1* and *Emx2*, which are key regulators of neurogenesis in the telencephalon^89, 90^. Likewise, Astro-CB cells express *Pax3*, which is also highly expressed in the CB GABAergic neurons. These observations are consistent with the notion that astrocytes and neurons are derived from common regionally distinct progenitors and share the common TFs for spatial patterning. Among other astrocyte-related subclasses, *Nkx2-2* is specific to Bergmann glia, *Rax* to tanycytes, *Myb* to ependymal cells, *Spdef* to hypendymal cells, and *Lef1* to CHOR. Some TFs are widely expressed but display specific expression patterns among related cell types. For example, *Tshz2* has much higher expression in Astro-OLF than in other astrocytes (**Extended Data Figure 8d)**.

The spatial distribution of all non-neuronal cell types in the mouse brain was confirmed and further refined by the MERFISH data. For example, we observed an inside-outside spatial gradient in MOB among the four OEG clusters (**Extended Data Figure 8g**). In addition to being widely distributed across the brain, oligodendrocytes are also highly concentrated in white-matter fiber tracts (**Extended Data Figure 8h-j)**. In contrast, the 1024 OPC NN_2 supertype is found mostly in gray-matter areas (**Extended Data Figure 8i**).

Of all the non-neuronal cell types, the Astro-Epen class exhibits the most diverse spatial patterns^91, 92^. Region-specific astrocytes Astro-OLF, Astro-TE, Astro-NT and Astro-CB are arranged in the UMAP in an anterior-to-posterior order (**Figure 4c**), consistent with their spatial patterning. Astro-TE cluster 5115, located in the lateral ventricle bordering rostral dorsal striatum, and clusters belonging to the Astro-OLF subclass (5119, 5120, 5118, 5116 and 5117) match the path of the rostral migratory stream (RMS)^93–95^. The trajectory of these astrocyte clusters on the UMAP matches well with the corresponding spatial gradients. Astro-TE cluster 5110 is located at the pia of telencephalon (**Figure 4c**) and has high expression of *Gfap* (**Extended Data Figure 8e**), consistent with the definition of interlaminar astrocytes (ILA)^96^.

Other clusters (5104, 5105, 5106, 5107) in the Astro-NT subclass are also localized at the pia with high expression of *Gfap*, which we hypothesize to be ILAs outside telencephalon. Besides *Gfap*, these clusters also have specific expression of *Atoh8* and *Myoc* (**Extended Data Figure 8e**). Other astrocyte-related subclasses, Astroependymal, Tanycyte, Hypendymal, Ependymal, and CHOR, line different parts of the ventricles throughout the brain (**Figure 4d**).

VLMC types^29, 97^ also show highly specific spatial and colocalization patterns. Clusters 5174, 5175, 5176 and 5177 are located at the pia, in contrast to clusters 5179 and 5178 which are scattered widely in the brain (**Figure 4e**). Interestingly, we found highly specific spatial colocalization between VLMC cluster 5181 and Tanycyte clusters (**Figure 4f**), between VLMC cluster 5180 and Ependymal/CHOR clusters (**Figure 4g**), and between pia specific VLMC clusters and ILAs (**Figure 4h**). Marker genes for VLMC clusters are enriched in extracellular matrix components and transmembrane transporters, including collagens and solute carriers with distinct cell type specificity (**Extended Data Figure 8f**). Interactions between various VLMC and astroependymal cell clusters, together with arachnoid barrier cells (ABC), likely regulate the movement of nutrients across the blood brain barrier^97^. The tanycyte-interacting VLMC cluster 5181 does not express many markers present in other VLMC types but has specific expression of transmembrane genes *Tenm4* and *Tmtc2*. This interaction may play roles in the brain-cerebrospinal fluid (CSF) barrier at the median eminence (ME)^98^.

Cell proliferation and neuronal differentiation continue in adulthood only in restricted areas of the brain^99^. The two main adult neurogenic niches are the dentate gyrus (DG) and the subventricular zone (SVZ) lining the lateral ventricles. The first gives rise to the excitatory DG granule cells, whereas the second produces migrating cells that follow the rostral migratory stream (RMS) and in the MOB differentiate into inhibitory granule cells^95, 100, 101^. We identified two subclasses of immature neurons, 278_MOB-STR-CTX Inh IMN and 279_DG-PIR Ex IMN, and grouped them with GABAergic neuron subclasses in MOB^102^ and glutamatergic granule cells in DG to form the MOB-DG-IMN class (**Table 1**, **Supplementary Table 7**, **Extended Data Figure 4**). We also uncovered relatedness between the Cajal-Retzius (CR) cells mostly found in HPF (subclass 280) and the MOB glutamatergic subclass (subclass 77) which are likely mitral and tufted cells^102^, and grouped them into the MOB-CR Glut class (**Table 1**, **Supplementary Table 7**, **Extended Data Figure 4**).

The scRNA-seq data show a trajectory from immature neurons to mature neurons in DG, and the MERFISH data corroborate that the immature neurons are located in the subgranular zone of DG, whereas the mature neurons reside in the dentate granular cell layer (**Figure 4i,j**). It seems, however, that the scRNA-seq data might not have captured all cell states along the maturation trajectory based on the gaps between clusters in the UMAP. Various studies have tried to capture the transitional states between neural stem and neuronal progenitor cells in the DG with most making use of transgenic mice to isolate specific states^103, 104^.

The migrating neurons in the RMS are separated from the parenchyma by astrocytes that form tunnels through which the cells migrate^94, 105^. RMS astrocytes (**Figure 4c,** cluster 5115 and the Astro-OLF subclass identified here) are molecularly distinct – they create a migration-permissive environment by providing soluble and non-soluble cues to the migrating neurons^93, 94^. In this well-orchestrated process, the neuroblasts, in turn, prevent astrocytic processes from invading the RMS by secreting *Slit1*, which acts on astrocytic *Robo* receptors to repel astrocytic processes out of the migratory path^94^. Our data showed two main cell populations arising from RMS into MOB; clusters that populate the inner granule and mitral cell layers (**Figure 4i,k**, trajectory 2), and clusters that populate the outer glomerular layer (**Figure 4i,l**, trajectory 3). Immature neurons in the SVZ and RMS are marked by the expression of cell cycle-associated genes like *Top2a* and *Mki67* (**Extended Data Figure 9**). As the MOB neurons exit the RMS, they express markers like *Sox11* and *S100a6* genes^106^, whereas the mature neurons in the MOB are marked by the expression of *Frmd7*.

### Transcription factor modules across the whole mouse brain

Transcription factors are considered key regulators of cell type identity^64, 65^. To evaluate the correspondence of TF expression to transcriptomic cell types, we calculated the number of differentially expressed (DE) TFs between each pair of divisions, classes, subclasses, or pairs of clusters within a subclass (**Figure 5a**). We then compared cross-validation accuracy of class, subclass and cell type recall using classifiers built based on all 8,108 DEGs, randomly selected 499 DEGs, or 499 TF marker genes (**Supplementary Table 8**, **Figure 5b**). The median cluster recall accuracy of cross-validation with TFs is between that of all DEGs and the random subset of DEGs. The cross-validation accuracy of subclass recall with TFs is 0.93, which is very similar to the accuracy with all DEGs (0.97), whereas the accuracy using the random subset of DEGs is much lower. The confusion matrix between the assigned and predicted subclasses in cross validation using classifiers trained on the 499 TF markers shows a high degree of concordance (**Figure 5c**). These results quantify the strong role of TFs in determining cell type identities.

We identified a large set of TF co-expression modules (**Methods**) that are selectively expressed in specific groups of cell types at all hierarchical levels and hence may define identities of these groups of cell types (**Figure 5d,e**, **Supplementary Table 8**). A pallium glutamatergic specific module includes *Tbr1* and *Satb2*. Immediate early genes *Egr3* and *Nr4a1* are also highly expressed in pallium glutamatergic neurons, while *Fos* and *Fosb* have more uniform expression. The bHLH transcription factors including *Neurod1*, *Neurod2*, *Neurod6* and *Bhlhe22* are widely expressed in many types of neurons but have highest expression in pallium glutamatergic cells. The *Dlx1*, *Dlx2*, *Dlx5*, *Dlx6*, *Arx*, *Sp8* and *Sp9* module is specific to GABAergic neurons in telencephalon, while the *Gata3*, *Gata2* and *Tal1* module is specific to GABAergic neurons in MB and pons. Interestingly, the latter gene module is best known as master regulator of hematopoietic development^107^, and is an example of re-purposing the same transcription factor module for specifying cell types in different systems. *Gbx2*, *Shox2* and *Tcf7l2* are highly expressed in thalamus glutamatergic neurons, while *Shox2* and *Tcf7l2* are also expressed in MB. *Hox* genes are specific to MY GABAergic and glutamatergic neurons. We also identified a TF module for the Astro-Epen cell class, including *Sox9*, *Gli2*, *Gli3*, and *Rfx4*, and several distinct modules for other non-neuronal cell subclasses.

For most other modules, each module consisted of a few TFs that are homologs, e.g., *Nfia/b/x*, the *Zic* family, the *Irx* family, the *Ebf* family, *En1/2*, *Lhx6/8*, *Six3/6*, and *Pou4f1/2/3*. Some of these homologs are located next to each other on the same chromosome, such as *Dlx1/2*, *Dlx5/6*, *Irx1/2*, *Irx3/5*, *Zic1/4*, *Zic2/5*, and *Hoxb2-8*. These homologs are likely located within the same chromatin domains, regulated by the same enhancers, and have highly similar expression patterns. Many co-expressed homologs show subtle but interesting distinctions. Consistent with the well-studied roles of *Hox* genes in regulating A-P Axis in development^108^, *Hoxb2/3* have broader expression than *Hoxb4/5*, and *Hoxb8* has the most restricted expression pattern in posterior lateral MY, in the order that is consistent with their locations on the chromosome.

While not very close on the chromosomes, *Nfia/b/x* regulate cell type differentiation in many tissues^109–111^, function as homo- or hetero-dimers, and bind to largely common targets^112^. Similar interactions between homologs have been reported for many other families, such as *Ebf*^113^ and *Irx*^114^. Finally, we identified a set of TFs such as *Meis1/2* and *Nr2f1/2* that are widely expressed but delineate neighboring subclasses and clusters and show local spatial gradients.

We further identified specific TFs that could define each node and branch in the dendrogram. Most subclasses could be uniquely specified by a combination of TFs located at all upper-level nodes, and some nodes and branches could be defined by one or just a few TFs (**Extended Data Figure 10a**). For example, transcription factor *Otp* marks several distinct populations in CNU, HY, MB and HB. We identified additional TFs to further distinguish these populations (**Extended Data Figure 10b**). *Otp*+ subclasses express *Foxg1* in CNU and anterior HY, *Ebf1*, *Ebf2*, *Irx2*, *En1* and *En2* in various MB areas, and *Pax3* in HB areas (**Extended Data Figure 10c**). Additional TFs provide finer separation. For example, subclasses 200/201 both express *En1/2*, but *Pax7* specifically in 201 and *Pax8* in 202. Both subclasses are located at ventral MB bordering pons, but 201 is more lateral, consistent with the spatial expression pattern of *Pax7* and *Pax8*. Similarly, subclasses 194/195 both express *Ebf1/2*, but *St18* specifically in 195 and *Evx2* in 194. Subclass 195 is located posteriorly at gigantocellular reticular nucleus (GRN) in MY while 194 is located more anteriorly at pontine reticular nucleus (PRNr) in pons. Together, these TF combinations delineate all the *Otp*+ subclasses.

While many TF homologs are co-expressed (**Figure 5e**), they can also show distinct expression patterns. We studied systematically the expression patterns of several TF families (**Extended Data Figure 11)**, including forkhead box (*Fox*), Krüppel-like factor (*Klf*), LIM Homeobox (*Lhx*), NKX-homeodomain (*Nkx*), Nuclear Receptors (*Nr*), Paired box (*Pax*), POU domain (*Pou*), Positive Regulatory Domain (*Prdm*), SRY-related HMG-box (*Sox*), and T-box (*Tbx*), all of which have been shown to play important roles in spatial patterning, cell type specification and differentiation during development^115–122^. In each family, only the TF markers identified in this study are included here. Members of the same TF family evolved from common ancestors, have strong sequence conservation, and very similar DNA binding motifs. Revealing their distinct cell type specificity provides deeper insights into the evolution of these TF families.

Particularly intriguing is the LIM Homeobox family, which can be split into multiple groups with complementary expression patterns that together cover most cell types in the brain. *Lhx2* and *Lhx9* are co-expressed in TH and MB glutamatergic types, but *Lhx2* is also specifically expressed in the pallium IT-ET types. *Lhx6* and *Lhx8* are co-expressed in some CNU/HY GABAergic types, but *Lhx6* is also specifically in MGE types. *Lhx1* and *Lhx5* are co-expressed in HY MM, MB and HB cell types, and much more highly in GABAergic than glutamatergic types. *Lmx1a* and *Lmx1b* are co-expressed in HB glutamatergic and MB dopaminergic cell types, but *Lmx1b* is also specifically in MB/HB serotonergic types. *Lhx3* and *Lhx4* are co-expressed in very specific glutamatergic types in pons and pineal gland. *Isl1* is widely expressed in HY/CNU, and much more highly in GABAergic than glutamatergic types. Interestingly, the grouping of *Lhx* members based on the gene expression patterns exactly matches their phylogeny tree based on their coding sequences^121^ and aligns with the sub-family definition.

### Brain region-specific cell type features

Characterizing the global features of cell type composition of regions across the brain complements our study of cell type diversity. We found that the numbers of cell types/clusters identified in different regions do not correlate with the numbers of cells profiled (**Figure 6a**). Rather, region-specific characteristics dominate. The hypothalamus, midbrain and hindbrain regions contain the largest numbers of clusters, indicating a high degree of cell type complexity, consistent with these regions having many small and heterogeneous subregions. Thalamus and cerebellum, on the other hand, contain the smallest numbers of clusters, suggesting lower complexity. Surprisingly, despite orders of magnitude more cells profiled in the pallium due to the many subregions contained within it (including isocortex, HPF, OLF and CTXsp, each containing multiple subregions) and its overall 4-15× larger volume compared to other major brain structures (**Supplementary Table 1**), we found an intermediate number of clusters for the entire pallium, similar to the other telencephalic structure, the subpallial CNU (**Figure 6a**).

The numbers of all genes or all transcription factors detected above a threshold per neuronal cluster are similar across all brain structures. However, when examining the homeobox TF gene family specifically, more homeobox TFs per neuronal cluster are detected in HB compared to all other structures, consistent with the unique roles these TFs play in hindbrain development^108^ (**Figure 6b**). We calculated the numbers of DEGs between each pair of clusters within a brain region, divided the numbers into nine quantiles based on similarities (i.e., higher similarity would yield fewer number of DEGs) and plotted their distribution by quantiles (**Figure 6c**).

Interestingly, we found that in regions with larger numbers of clusters, i.e., HY, MB and HB, their clusters are more similar to each other within each region, suggesting that cell types in these regions have lower diversity and are less hierarchical. In contrast, in regions with smaller numbers of clusters, i.e., CB, TH and Pall, there are wide differences in similarities between cell types, thus, cell types in these regions may be more diverse and hierarchical. CNU exhibits an intermediate level of diversity. The results show that HY, MB and HB have more numerous cell types, but the cell types are more like each other. We also calculated the 3D spatial span of each cluster based on the MERFISH dataset and aggregated the spans of all clusters within each brain region (**Figure 6d**). Each region shows its own unique characteristics, with clusters in pallium having much larger spans suggesting sharing across subregions, and clusters in HY having much smaller spans suggesting more restricted localization.

In addition to regional specificity of cell types, we also observed continuity across major brain regions by identifying a specific set of cell types that are shared or transitioning between brain regions (**Figure 6e-g**). For example, cells belonging to glutamatergic subclasses 12, 98 and 99 are found in both pallial OLF (e.g., cortical amygdala area, COA) and subpallial sAMY (e.g., medial amygdala nucleus, MEA) regions. The GABAergic subclass 46 contains neurons in both isocortex (the Sst Chodl cells^25, 28^) and dorsal and ventral striatum (STRd and STRv). Subclass 54 is shared between STR and PAL. Subclass 56 is a transitional type between LSX and PAL. Subclasses 86 and 92 are shared between sAMY and PAL. GABAergic subclasses 102-105 and glutamatergic subclasses 128 and 129 are shared between PAL and anterior HY. Subclass 149 is transitional between HY and MB. Glutamatergic subclasses 158 and 159 and GABAergic subclasses 213 and 214 are transitional between TH and MB. Glutamatergic subclasses 153 and 154 and GABAergic subclasses 201, 209, 210, 223 and 237 are transitional between MB and pons.

We investigated sex differences in the whole mouse brain transcriptomic cell type atlas. We identified 28 clusters across 13 subclasses with a skewed distribution of cells derived from the two sexes (**Figure 1a**, **Supplementary Table 7**). Of these, 5 are small, sex-specific clusters: clusters 211, 1402, 2536 and 2538 are male-specific and cluster 2207 is female-specific. The 23 sex-dominant clusters include 1404, 2058, 2062, 2065, 2088, 2089, 2154, 2196, 2204 and 3612, which contain mostly cells from female donors, and clusters 1396, 1407, 1409, 1781, 1843, 2048, 2057, 2061, 2150, 2195, 3359, 3716 and 3952, which contain mostly cells from male donors. Based on the MERFISH data, these clusters mostly reside in specific regions of PAL, sAMY, HY and HB.

Within the whole mouse brain scRNA-seq dataset, we also collected a complete subset of data covering all brain regions from the dark phase of the circadian cycle (**Supplementary Table 2**, total 1,121,542 10xv3 cells). All the dark-phase transcriptomes were included in the overall clustering analysis. In all but one subclass, they are found commingled with the corresponding light-phase transcriptomes (the exception being subclass 253, with only 22 cells that are all from the light phase) (**Extended Data Figure 3**, **Supplementary Table 7**). Out of all 5,200 clusters, there are 271 clusters that do not contain dark-phase cells, while none contain dark-phase cells only. Detailed gene expression analysis at class and subclass levels revealed widespread expression differences of canonical circadian clock genes between the light and dark phases (**Extended Data Figure 12**). Across many neuronal and non-neuronal classes and subclasses throughout the brain, nearly all clock genes show consistently higher expression levels in the dark phase than the light phase, except for *Arntl* which displays an opposite pattern.

Interestingly, the 283_Pineal Crx Glut subclass, which is found located in the dorsal part of the third ventricle and on top of superior colliculus (SC) in the MERFISH data and likely represents the pinealocytes that evolved from photoreceptor cells and secret melatonin^123^, has particularly robust circadian gene expression fluctuations (**Extended Data Figure 12b,c**). Furthermore, in the 82_SCH Gaba subclass, which is specific to the suprachiasmatic nucleus (SCH), the circadian pacemaker of the brain, most clock genes (e.g., *Per1*, *Per3*, *Dbp*, *Nr1d1*, *Nr1d2*) have higher levels of expression in the light phase than the dark phase, suggesting that the pacemaker cells are at a different phase of the circadian cycle of gene expression from the rest of the brain, consistent with previous findings^124^ (**Extended Data Figure 12b,c**). Intriguingly, the vascular 297_ABC NN subclass also displays a similar phase shift. These results suggest that our whole mouse brain transcriptomic cell type atlas also captured circadian state-dependent gene expression changes. While supervised analysis can reveal these changes, our cell type classification is not significantly affected by the different circadian states.

## DISCUSSION

In this study, we created a comprehensive, high-resolution transcriptomic cell type atlas for the whole adult mouse brain based on the combination of two whole-brain-scale datasets: a scRNA-seq dataset of ∼7 million cells and a MERFISH dataset of a similar scale (∼4.3 million cells from the AIBS MERFISH brain). We used ∼4.1 million high-quality single-cell transcriptomes after stringent QC to create a transcriptomic cell type taxonomy. We used the MERFISH data, which were generated using marker genes derived from the whole-brain transcriptomic taxonomy, to annotate the spatial location of each subclass and each cluster in the taxonomy. We then built a hierarchically organized transcriptomic and spatial cell type atlas with five nested levels: 7 divisions, 32 classes, 306 subclasses, 1,045 supertypes and 5,200 clusters (**Figure 1**). The neuronal cell type composition in each major brain structure were systematically analyzed (**Figure 2**) and distinct features in different brain structures identified (**Figure 6**). We discovered many sets of neuronal types with varying degrees of similarity with each other, including transitional neuronal types across regions as well as highly distinct neuronal types. We also systematically analyzed all divisions of non-neuronal cell types as well as immature neuronal types present in the adult brain, and identified their unique spatial distribution and spatial interaction patterns (**Figure 4**). Finally, we systematically characterized cell-type specific neurotransmitters, neuropeptides, and transcription factors, and discovered unique characteristics for each as discussed below. This large-scale study allowed us to delineate several principles regarding cell type organization across the whole mouse brain. It provides a benchmark reference cell type atlas as a resource for the community that will enable many more discoveries in the future.

One of the most striking findings from our study is the high degree of correspondence between transcriptomic identity and spatial specificity. Every subclass (and all supertypes and many clusters within each) has a unique and specific spatial localization pattern within the brain.

Furthermore, the relative relatedness between transcriptomic types (as revealed in 2D and 3D UMAPs) is strongly correlated with the spatial relationship between them. Transcriptomically related cell types are often found in the same region, or in some cases in related regions that have a common developmental origin. Transitioning cell types in the transcriptomic space are also found crossing regional boundaries. We believe that the strong correspondence between transcriptomic and spatial specificity and relatedness indicates the importance of anatomic specialization of cell types and lends strong support to the robustness and validity of our transcriptomics-based cell type classification. Given that spatial organization of the brain is laid out during development, we further hypothesize that developmental origins and relationships may be inferred from the adult stage transcriptomic profiles of the cell types.

Another striking finding is the distinct features of cell type organization between the major brain structures (**Figure 6**). The anterior and dorsal brain structures, including olfactory areas, isocortex, hippocampal formation, dorsal striatum, thalamus, and cerebellum, contain cell classes and types that are highly distinct from the other parts of the brain. Cell types in these structures also tend to be more widely distributed, often shared between neighboring regions or subregions. In contrast, cells from the ventral part of the brain, including striatum-like amygdala nuclei, ventral pallidum, hypothalamus, midbrain, pons and medulla, form numerous small clusters that are closely related to each other. And these cell types often have restricted spatial localization, forming the small nuclei characteristic of these regions. This dichotomy between the roughly dorsal and ventral parts of the brain may reflect the different evolutionary histories of these brain structures.

There are several remarkable differences between neuronal and non-neuronal cell types. While neuronal types constitute the vast majority of cell types in the brain and exhibit high regional specificity, non-neuronal types are generally more widely distributed, except for astrocytes which have multiple subclasses with regional specificity. However, even for those widely distributed non-neuronal types, at the cluster level we observed a great degree of spatial specificity, especially for astrocytes, ependymal cells, tanycytes and VLMCs, indicating specific neuron-glia and glia-vasculature interactions (**Figure 4a-h**). We also identified several groups of immature neuronal types and could infer their trajectories to mature neuronal types in olfactory bulb and dentate gyrus based on their spatial localization and transitioning gene signatures (**Figure 4i-l**).

As example case studies, we examined the discovered diversity in neurotransmitter and neuropeptide expression in cell types across the brain. We found a diverse set of neuronal clusters with glutamate-GABA dual transmitters from many brain regions (**Figure 3a-d**). We identified all cell types expressing different modulatory neurotransmitters and found that they often co-release glutamate and/or GABA. Intriguingly, the neuromodulatory cell types are usually not completely segregated from other neuronal types, but often have closely related glutamatergic and/or GABAergic clusters within the same subclass, showing a high degree of heterogeneity in neurotransmitter content in these cell populations (**Figure 3e-h**). Our assignment of neurotransmitter types based on the most specific transporter genes is conservative; there may be even more diversity in neurotransmitter co-release patterns if alternative transmitter release routes are considered^76, 79, 80^. Similarly, there is a wide spectrum of expression patterns among the different neuropeptide genes, some widely expressed in many cell types while others highly specific to one or few cell types (**Extended Data Figure 7**).

Furthermore, there are myriad co-expression combinations of two or more neuropeptide genes in many neuronal clusters (**Supplementary Table 7**). These results support the extraordinary diversity in intercellular communications in the brain.

We found that transcription factors are highly predictive in determining cell type classification. Transcription factors are known to play major roles in patterning brain regions, defining neural progenitor domains and specifying cell type identities during development. Here, we found that in the adult mouse brain, transcription factors also are major determinants in defining cell types across all regions of the brain. Out of the 8,108 marker genes we identified for the 5,200 cell clusters, 499 are TF genes. In cross-validation tests, the 499 TF genes predicted cell subclass and cluster identities nearly as well as all the marker genes together (**Figure 5a-c**). A hierarchical tree derived from hierarchical clustering using the 499 TF genes alone better recapitulated the existing knowledge about cell types and their spatial relationships at class and subclass levels than using all the marker genes, and thus we used the TF-derived tree to represent the cell type taxonomy (**Figure 1a**). We identified TF genes and co-expression modules specific to top hierarchical levels and most branches of the cell type taxonomy (**Figure 5e, Extended Data Figure 10, 11**). We also found several different modes of coordination among TFs. The first mode is the coordinated expression of different TFs (often pairs of TFs) within the same TF gene family in specific cell types. The second is the combination of TFs at different hierarchical branch levels to collectively define the identity of the leaf-node subclasses. The third represents the intersection between different sets of TFs that define molecular identity or spatial specificity, respectively, within a cell type. These findings reveal how transcription factors form the combinatorial code to lay out the highly complex cell type landscape.

We must also emphasize the great computational challenges in analyzing these large and highly complex datasets and the two main caveats for the results presented here. First, due to the difficulty in dissociating and isolating intact cells from the adult brain tissue, especially in highly myelinated areas, our scRNA-seq dataset contains many kinds of low-quality cells, including damaged cells, debris, doublets or mixed debris of various cell type combinations. These low-quality transcriptomes could be mistaken for real cell types, part of a cell type continuum, or transitional cell types in clustering results. They could also lead to wrong mapping of MERFISH cells as we discovered in our analysis. To generate a high-quality transcriptomic cell type atlas with precise spatial annotation, we developed a set of QC metrics that are more stringent than those widely used in the field and therefore we failed a high proportion of cells from our scRNA-seq dataset (**Extended Data Figure 1**). During this process, it is likely that some cell types were more selectively depleted than others, especially large neurons that are more vulnerable to damage during tissue dissociation, e.g., Purkinje cells and large motor neurons in the midbrain and hindbrain. Thus, cell types in the midbrain and hindbrain may not be fully represented or fully resolved in our whole mouse brain transcriptomic cell type atlas. We observed that many of the QC-failed transcriptomes resemble single-nucleus transcriptomes; they might be still useful for specific analysis purposes and could be rescued from our dataset to recover certain cell types in the future. These observations highlight the importance of collecting very large multimodal datasets in constructing cell type atlases that are complete, accurate, and permanent.

Second, although we only used the selected high-quality single-cell transcriptomes to construct the cell atlas, the relationships between the large number of cell types across the entire brain are still highly complex and impossible to be fully captured by a one-dimensional hierarchical tree or two-dimensional UMAPs. The transcriptomic profile of each cell is multi-dimensional, containing not only information about the cell type identity, but also information about many other aspects of the cellular properties such as spatial location, connectivity, function, or a particular cell state. We conducted extensive iterative clustering to resolve all dimensions of variation at the cluster level. Thus, not every cluster may represent a true cell type; our categorization scheme may not be perfectly reflecting the brain-wide cell type organization either and will need to be revised in the future with better computational methods and/or more experimental evidence (especially developmental data). Finally, due to the sheer scale of the atlas, we have not extensively searched and utilized the vast amount of existing data and knowledge about cell types in many parts of the brain to help better annotate our cell type atlas. Moving forward, it will be critical to engage the neuroscience community to collectively annotate, refine and enhance this whole mouse brain cell type atlas, and an online platform to facilitate this will be needed.

In conclusion, the whole-brain transcriptomic and spatial cell type atlas establishes a foundation for deep and integrative investigations of cell type and circuit function, development, and evolution of the brain, akin to the reference genomes for studying gene function and genomic evolution. The atlas provides baseline gene expression patterns that allow investigation of the dynamic changes in gene expression and cellular function in different physiological and diseased conditions. It enables creation of cell type-targeting tools for labeling and manipulating specific cell types to probe and modify their functions *in vivo*. The atlas provides a foundational framework for organizing and integrating the vast knowledge about the brain structure and function, facilitating the extraction of new principles from the extraordinarily complex cell type and circuit landscape. It provides a starting point for generating similarly comprehensive and detailed cell type atlases for other species as well as across developmental times, enabling cross-species comparative studies and gaining mechanistic insights on the genesis of cell types and circuits in the mammalian brain. Understanding the conservation and divergence of cell types between human and model organisms will have profound implications for the study of human brain function and diseases.

## METHODS

### Mouse breeding and husbandry

All procedures were carried out in accordance with Institutional Animal Care and Use Committee protocols at the Allen Institute for Brain Science. Mice were provided food and water *ad libitum* and were maintained on a regular 14:10 hour day/night cycle at no more than five adult animals of the same sex per cage. Mice were maintained on the C57BL/6J background. We excluded any mice with anophthalmia or microphthalmia.

We used 95 mice (41 female, 54 male) to collect 2,492,084 cells for 10xv2 and 222 mice (112 female, 110 male) to collect 4,466,283 cells for 10xv3. Animals were euthanized at P53-59 (*n* = 141), P50-52 (*n* = 3), or P60-71 (*n* = 173). No statistical methods were used to predetermine sample size. All donor animals used in this study are listed in **Supplementary Table 2.**

Transgenic driver lines were used for fluorescence-positive cell isolation by FACS to enrich for neurons. Most cells were isolated from the pan-neuronal *Snap25-IRES2-Cre* line crossed to the *Ai14*-tdTomato reporter^125, 126^ (279 out of 317 donors) (**Supplementary Table 2**). A small number of *Gad2-IRES-Cre/wt;Ai14/wt* (6 donors) and *Slc32a1-IRES-Cre/wt;Ai14/wt* mice (4 donors) were used for fluorescence-positive cell isolation to enrich for the sampling of GABAergic neurons in HIP, OLF and CB. For unbiased sampling without FACS, we used either *Snap25-IRES2-Cre/wt;Ai14/wt or Ai14/wt* mice.

The number of mice contributing to each cluster varies between 2 and 266, with an average of 19 and median of 14. There are 19 clusters that have fewer than 4 donor animals each. Thus, individual mouse variability should not affect cell type identities (**Extended Data Figure 3**).

For cell collection during the dark phase of the circadian cycle, mice were randomly assigned to circadian time groups at time of weaning and housed on the reversed 12:12 hour light/dark cycle. Brain dissections for all groups took place in the morning. From 267 donors, 5,836,825 cells were collected during the light phase of the light-dark cycle. For 50 donors, 1,121,542 cells across the whole brain were collected during the dark phase of the light-dark cycle (**Supplementary Table 2**).

### Single-cell RNA-sequencing Single-cell isolation

We used the Allen Mouse Brain Common Coordinate Framework version 3 (CCFv3; RRID: SCR_002978) ontology^63^ (http://atlas.brain-map.org/, **Supplementary Table 1**) to define brain regions for profiling and boundaries for dissection. We covered all regions of the brain using sampling at top-ontology level with judicious joining of neighboring regions (**Supplementary Table 3, Extended Data Figure 1d-e**). These choices were guided by the fact that microdissections of small regions were difficult. Therefore, joint dissection of neighboring regions was sometimes necessary to obtain sufficient numbers of cells for profiling.

Single cells were isolated by adapting previously described procedures^28, 127^. The brain was dissected, submerged in ACSF, embedded in 2% agarose, and sliced into 350-μm coronal sections on a compresstome (Precisionary Instruments). Block-face images were captured during slicing. Regions of interest (ROIs) were then microdissected from the slices and dissociated into single cells as previously described^28^. Fluorescent images of each slice before and after ROI dissection were taken at the dissection microscope. These images were used to document the precise location of the ROIs using annotated coronal plates of CCFv3 as reference.

Dissected tissue pieces were digested with 30 U/ml papain (Worthington PAP2) in ACSF for 30 minutes at 30°C. Due to the short incubation period in a dry oven, we set the oven temperature to 35°C to compensate for the indirect heat exchange, with a target solution temperature of 30°C. Enzymatic digestion was quenched by exchanging the papain solution three times with quenching buffer (ACSF with 1% FBS and 0.2% BSA). Samples were incubated on ice for 5 minutes before trituration. The tissue pieces in the quenching buffer were triturated through a fire-polished pipette with 600-µm diameter opening approximately 20 times. The tissue pieces were allowed to settle and the supernatant, which now contained suspended single cells, was transferred to a new tube. Fresh quenching buffer was added to the settled tissue pieces, and trituration and supernatant transfer were repeated using 300-µm and 150-µm fire polished pipettes. The single cell suspension was passed through a 70-µm filter into a 15-ml conical tube with 500 µl of high BSA buffer (ACSF with 1% FBS and 1% BSA) at the bottom to help cushion the cells during centrifugation at 100 x g in a swinging bucket centrifuge for 10 minutes. The supernatant was discarded, and the cell pellet was resuspended in the quenching buffer. We collected 1,508,284 cells without performing FACS. The concentration of the resuspended cells was quantified, and cells were immediately loaded onto the 10x Genomics Chromium controller.

To enrich for neurons or live cells, cells were collected by fluorescence-activated cell sorting (FACS, BD Aria II) using a 130-μm nozzle. Cells were prepared for sorting by passing the suspension through a 70-µm filter and adding Hoechst or DAPI (to a final concentration of 2 ng/ml). Sorting strategy was as previously described^28^, with most cells collected using the tdTomato-positive label. 30,000 cells were sorted within 10 minutes into a tube containing 500 µl of quenching buffer. We found that sorting more cells into one tube diluted the ACSF in the collection buffer, causing cell death. We also observed decreased cell viability for longer sorts. Each aliquot of sorted 30,000 cells was gently layered on top of 200 µl of high BSA buffer and immediately centrifuged at 230 x g for 10 minutes in a centrifuge with a swinging bucket rotor (the high BSA buffer at the bottom of the tube slows down the cells as they reach the bottom, minimizing cell death). No pellet could be seen with this small number of cells, so we removed the supernatant and left behind 35 µl of buffer, in which we resuspended the cells. Immediate centrifugation and resuspension allowed the cells to be temporarily stored in a high BSA buffer with minimal ACSF dilution. The resuspended cells were stored at 4°C until all samples were collected, usually within 30 minutes. Samples from the same ROI were pooled, cell concentration quantified, and immediately loaded onto the 10x Genomics Chromium controller.

### cDNA amplification and library construction

For 10x v2 processing, we used Chromium Single Cell 3’ Reagent Kit v2 (120237, 10x Genomics). We followed the manufacturer’s instructions for cell capture, barcoding, reverse transcription, cDNA amplification, and library construction^128^. We targeted sequencing depth of 60,000 reads per cell; the actual average achieved was 54,379 ± 34,845 (mean ± SD) reads per cell across 299 libraries.

For 10x v3 processing, we used the Chromium Single Cell 3′ Reagent Kit v3 (1000075, 10x Genomics). We followed the manufacturer’s instructions for cell capture, barcoding, reverse transcription, cDNA amplification and library construction^129^. We targeted a sequencing depth of 120,000 reads per cell; the actual average achieved was 83,190 ± 85,142 reads per cell across 482 libraries.

### Sequencing data processing and QC

Processing of 10x Genomics libraries was performed as described previously^28^. Briefly, libraries were sequenced on the Illumina NovaSeq6000, and sequencing reads were aligned to the mouse reference transcriptome (M21, GRCm38.p6) using the 10x Genomics CellRanger pipeline (version 6.1.1) with default parameters.

To remove low quality cells, we developed a stringent QC process. Cells were first classified into broad cell classes after mapping to an existing, preliminary version of taxonomy, and cell quality was assessed based on gene detection, qc score, and doublet score. The qc score was calculated by summing the log transformed expression of a set of genes whose expression level is decreased significantly in poor quality cells. These are housekeeping genes that are strongly expressed in nearly all cells with a very tight co-expression pattern that is anti-correlated with the nucleus localized gene *Malat1* (**Supplementary Table 4)**. Out of the 62 such genes chosen, 30 are annotated as mitochondrial inner membrane category based on GO ontology cellular component, although they are not located on the mitochondrial chromosome. Some evidence suggests the mRNAs of some of these genes or their homologs are translocated to the mitochondrial surface^130, 131^. We used this qc score to quantify the integrity of cytoplasmic mRNA content, which tended to show bimodal distribution. Cells at the low end were very similar to single nuclei, which we removed for downstream analysis. Doublets were identified using a modified version of the DoubletFinder algorithm^132^ and removed when doublet score > 0.3. Using thresholds that were tailored to different cell classes, we filtered out 43% and 29% of cells and kept 2,546,319 cells and 1,769,304 cells for 10xv3 and 10xv2 data, respectively (**Extended Data Figure 1**). Threshold parameters and number of cells filtered are summarized in **Supplementary Table 4.**

### Clustering single cell RNA-seq data

Clustering for both 10xv2 and 10xv3 datasets was performed independently using the in-house developed R package **scrattch.bigcat** (available via github https://github.com/AllenInstitute/scrattch.bigcat), which is a scaled-up version of R package scrattch.hicat^25, 28^ to deal with the increased size of datasets. Scrattch.bigcat adopted the parquet file format for storing sparse matrix, which allows for manipulation of matrices that are too large to fit in memory through memory mapping to files on disk. The whole gene count matrices were chunked to smaller parquet files with bin size of 50,000 for cells, and 500 for genes, which could be loaded efficiently and concurrently using the arrow package (https://github.com/apache/arrow/, https://arrow.apache.org/docs/r/).

We provide utility functions to convert and concatenate sparse matrices in R to this format, and functions for conversion between this format and other commonly used file formats such as h5, h5ad and Zarr. We also provide a function that loads any sub-matrix into the memory given the cell IDs and gene IDs. The choice of parquet format is based on its great performance in R, which allows continual usage of our legacy codebase. The major functions of scrattch.hicat package were rewritten and made available in scrattch.bigcat. We used the automatic iterative clustering method, iter_clust_big, which performed clustering in top down manner into cell types of increasingly finer resolution without any human intervention, while ensuring that all pairs of clusters, even at the finest level, were separable by stringent differential gene expression criteria as follows: for 10v2, q1.th = 0.4, q.diff.th = 0.7, de.score.th = 150, min.cells = 10; for 10xv3, q1.th = 0.5, q.diff.th = 0.7, de.score.th = 150, min.cells = 4. These criteria translated to at least 8 binary DEGs between any pair of clusters (each DEG’s contribution to de.score was capped at 20, so at least 8 genes were needed to exceed de.score.th of 150). Binary DEGs were defined as genes expressed in at least 40% cells in the foreground cluster in 10xv2, and 50% in 10xv3 (q1.th parameter), log2FC > 1, adj Pval < 0.01, and difference between the fraction of cells expressing the gene in foreground and background divided by the foreground fraction was greater than 0.7 (q.diff.th parameter).

To enhance scalability, a randomly subsampled set of cells to be clustered were loaded into memory to compute high variance genes and perform PCA, then projected to all the cells to obtain their reduced dimensions. Then Jaccard-Leiden clustering proceeded as before^28^.

### Differential gene expression analysis

We performed differential gene expression both at the clustering step for each iteration, and after clustering between all pairs of clusters. In our original scrattch.hicat package, we applied limma package^133^ to perform this analysis. Given the significant increase of data size and complexities of the taxonomy, we re-implemented this method that provides essentially identical results, but drastically improves performance and scalability. The method first scanned the whole log transformed cell-by-gene matrix once to compute, for each cluster and each gene, the average expression, the fraction of cells expressing the gene, and the sum of square of gene expression of all the cells within the cluster. These cluster level summary statistics were then used in the linear model equivalent to the one used in limma to compute the pvalue, adjusted pvalue, log fold change, and the contrast between foreground and background based on the fraction of cells expressing the gene. This process was massively parallelized. Clusters were grouped into bins, and the DEG analysis results were stored on disk in chunked parquet files, split based on which bin the foreground and background clusters belonged to. In this way, we were able to compute DEGs between ∼13.5 million pairs of clusters within a day on a single Linux server. Using the arrow package, we were able to query DEGs between any pairs of clusters very efficiently.

### Excluding noise clusters

Before proceeding with integration between 10xv2 and 10xv3 datasets, we first needed to remove noise clusters. The presence of such clusters can confuse the integration algorithm and reduce the cell type resolution. There are two main categories of noise clusters: clusters with significantly lower gene detection due to extensive drop out, and clusters due to doublets or contamination.

We first identified doublet clusters based on the co-expression of any pair of broad class marker genes using find_doublet_by_marker function in scrattch.bigcat package. To identify other doublet clusters, we searched for triplets of clusters A, B and C, wherein A was the putative doublet cluster, such that up-regulated genes of A relative to B largely overlapped with up- regulated genes in C relative to B, and up-regulated genes in A relative to C largely overlapped with up-regulated genes of B relative to C. This criterion ensured that A included the most distinguished signature of B and C. To rule out the possibility that A was a transitional type between B and C, we required that B and C could not be closely related types based on the correlation of their average gene expression of marker genes. After we systematically produced the list of all the candidate triplet clusters, the final determination was an iterative process that involved setting different thresholds and manual inspection of borderline cases.

After removing all doublet clusters, we then identified clusters with lower gene detection. To do that, we identified pairs of clusters such that one cluster with at least 50% fewer UMIs or >100 lower QC score, smaller size, and no more than one up-regulated gene relative to another cluster was identified as the low-quality cluster. In these cases, one cluster was a degraded version of another cluster and therefore removed.

We identified 933 noise clusters with 153,598 cells in 10xv3, and 201 noise clusters with 38,073 cells in 10xv2. 10xv3 noise clusters were removed from integration analysis but 10xv2 noise clusters were included accidentally. Fortunately, most of the cells from 10xv2 noise clusters were excluded in further QC steps after integration.

### Joint clustering 10xv2 and 10xv3 datasets

To provide one consensus cell type taxonomy based on both 10xv2 and 10xv3 datasets of ∼2M cells each, we scaled up the integrative clustering method^28^ and made it available via scrattch.bigcat package which extends the clustering pipeline described above to integrate datasets collected by different transcriptomic platforms. Analysis was performed as described before^28^ with minor modifications. To build the common graph that incorporates samples from all the datasets, both 10xv2 and 10xv3 were used as the reference datasets. The key steps in the pipeline are: 1) select anchor cells for each reference dataset, 2) select high variance genes in each reference dataset, prioritizing shared high variance genes, 3) compute K nearest neighbors (KNN) both within modality and cross modality, 4) compute Jaccard similarity based on shared neighbors, 5) perform Leiden clustering based on Jaccard similarity, 6) merge clusters based on total number and significance of conserved DEGs across modality between similar cell types, 7) repeat steps 1–6 for cells within a cluster to gain finer-resolution clusters until no clusters can be found, 8) concatenate all the clusters from all the iterative clustering steps and perform final merging as in step 6. For step 6, if one cluster had fewer than the minimal number of cells in a dataset (4 cells for 10xv3 and 10 cells for 10xv2), then this dataset was not used for DE gene computation for all pairs involving the given cluster. This step allows detection of unique clusters only present in some data types.

Compared to the previous version, the key improvement is step 3 for computing KNN. We used BiocNeighbor package (https://github.com/LTLA/BiocNeighbors) for computing KNN using Euclidean distance within modality and Cosine distance across modality using the Annoy algorithm (https://github.com/spotify/annoy). The Annoy index was built based on anchor cells for the reference dataset, and KNNs were computed in parallel for all the query cells. Due to significantly increased dataset sizes, the Jaccard similarity graph can be extremely large, impossible to fit in memory. The method down-samples the datasets based on a user specified parameter, and if the cluster membership of each modality is provided as input for integration algorithm, we down-sample cells by within-modality clusters, ensuring preservation of rare cell types. All the anchor cells were added to the down-sampled datasets. The Jaccard-Leiden clustering was performed on the down-sampled datasets, and the cluster membership of other cells were imputed based on KNNs computed in step 3.

The integration algorithm generated 5,283 clusters, which were used to build cell type taxonomy. During this process, additional noise clusters were identified by manual inspection, which exhibited abnormal QC statistics, abnormal expression of canonical markers, or absence in

MERFISH dataset. Most of these clusters were very small, likely doublets of damaged cells. After removing these additional noise clusters, the final taxonomy had 5,200 clusters with 4,058,049 cells.

### Marker gene selection

For each pair of clusters, we computed conserved DEGs (at least significant in one dataset, and at least 2-fold change in the same direction in the other datasets). We selected the top 15 DEGs in each direction and pooled such genes from all pairwise comparisons to generate a total of 8,108 gene markers (**Supplementary Table 5**).

### Assessing concordance of joint clustering between 10xv2 and 10xv3

We first compared the joint clustering result with the independent clustering result from each dataset. We then calculated the cluster means of marker genes for each dataset. For each marker gene, we computed the Pearson correlation between its average expression for each cluster across two different datasets to quantify the consistency of its expression at the cluster level between datasets (**Extended Data Figure 5d**). We performed a similar analysis between 10xv3 and MERFISH datasets.

### Imputation

To facilitate direct comparisons, we projected gene expression of the 10xv2 dataset to the 10xv3 dataset using the impute_knn_global function in the scrattch.bigcat package^28^. To achieve this, we leveraged the KNN matrices computed iteratively at each level of the cell type hierarchy.

During each iteration of the joint clustering, we used the average gene expression of the K nearest neighbors among the 10xv3 anchor cells as the imputed expression for each 10xv2 cell. At the top-level clustering, we imputed the expression for all genes. For each following iteration, we only imputed the expression of the DEGs computed for the cells involved in the given iteration. We used this iterative approach for imputation because the nearest neighbors, based on the genes chosen at the top level, may not reflect the distinction between the finer types, and the imputed values for the DEGs that define the finer types consequently are not accurate based on these nearest neighbors. Therefore, we deferred imputation of the DEGs between the finer types to the iteration when these types were defined. The key improvement of this function is parallelization of KNN computation and storing the output imputed matrix as file backed matrix (FBM) for scalability.

### UMAP projection

We performed PCA based on the imputed gene expression matrix of 8,108 marker genes using the 10xv3 reference. We down-sampled up to 100 cells per cluster, and further down-sampled up to 250K cells if the total exceeded this number, so that PCA could proceed without any memory issues. Again, the PCs based on sampled cells were projected to the whole datasets. We selected the top 100 PCs, then removing one PC with more than 0.7 correlation with the technical bias vector, defined as log2(gene count) for each cell. We used the remaining PCs as input to create 2D and 3D UMAPs^134^, using parameters nn.neighbors = 25 and md = 0.4. To prevent some of the big clusters taking up too much space, we down sampled up to 1000 cells per cluster to build the UMAP and impute the UMAP coordinates of the other cells based on KNN neighbors among the sampled cells in the PCA space.

### Building cell type hierarchy

To make the cell type complexity tractable at each level, we organized the 5,200 clusters into a hierarchy with 5 levels: division, class, subclass, supertype and cluster. After clusters were computed as descripted in **Joint clustering** section, we first defined subclasses by clustering the clusters. This was performed by Jaccard-Leiden clustering using the average expression of 499 TF marker genes of all the cells in each cluster, using 5 K nearest neighbors, and varying the resolution index of Leiden algorithm at 0.1, 0.2, 1, 5, and 8. We tried clustering using either all 8,108 marker genes or 499 TF marker genes only, and found the result based on TF marker list recapitulate existing knowledge of cell types including spatial distribution and lineage relationships better. The Leiden algorithm generated 32 groups at resolution index 0.2, which generated the initial version of “classes”, and 195 groups at resolution index 8, which generated the initial version of “subclasses”.

The initial fully automatically generated versions of classes and subclasses were visualized together with all the other metadata on UMAPs and on MERFISH sections using the single-cell data visualization tool cirrocumulus (https://cirrocumulus.readthedocs.io/en/latest/) for manual examination. We finetuned the borderline cases, and further split or merged some putative subclasses to reach the final definition of subclasses. We applied a similar process to define classes, and to achieve strict hierarchy, assigned all the clusters in one subclass to the same class. The classes were then grouped into divisions, informed by prior knowledge and the subclass taxonomy tree (see below). Finally, we applied the same Jaccard-Leiden algorithm to all the clusters within each subclass separately to define supertypes, using the union of the top 20 DEGs between all pairs of clusters within the subclass as features. Again, they were adjusted based on manual inspection of UMAPs and MERFISH sections after visualization on cirrocumulus to increase the consistency of supertype definitions between subclasses.

### Building subclass taxonomy tree

We built the subclass taxonomy tree using the average expression of 499 TF marker genes at subclass level, using the build_dend function in the scrattch.bigcat package as described previously^28^. Branches with length < 0.01 were removed from the tree, and the children of any removed node were re-assigned as children of the parent of the removed node. The tree captured relationships between closely related subclasses, but the hierarchy is not fully consistent with the “class” definition, as hierarchical clustering is not capable of capturing the continuous variations in multi-dimensional space. We used the cell type hierarchy, the taxonomy tree, the 2D and 3D UMAPs and the constellation plot all together to understand the overall cell type landscape and relationships between cell types.

### Constellation plot

The global relatedness between cell types was visualized using a constellation plot (**Extended Data Figure 4**). To generate the constellation plot, each transcriptomic subclass was represented by a node (circle), whose surface area reflected the number of cells within the subclass in log scale. The position of nodes was based on the centroid positions of the corresponding subclasses in UMAP coordinates. The relationships between nodes were indicated by edges that were calculated as follows. For each cell, 15 nearest neighbors in reduced dimension space were determined and summarized by subclass. For each subclass, we then calculated the fraction of nearest neighbors that were assigned to other subclasses. The edges connected two nodes in which at least one of the nodes had > 5% of nearest neighbors in the connecting node. The width of the edge at the node reflected the fraction of nearest neighbors that were assigned to the connecting node and was scaled to node size. For all nodes in the plot, we then determined the maximum fraction of “outside” neighbors and set this as edge width = 100% of node width. The function for creating these plots, plot_constellation, is included in scrattch.bigcat.

### Defining neighborhoods

We identified highly prevalent transitions between cell types at almost all levels. To study these transitions not captured by the strict hierarchical 5-level taxonomy, we defined multiple overlapping neighborhoods. For example, the transition cell types between CNU and Pallium Glut cell types were included in both Pallium Glut neighborhood and PAL-sAMY-HY neighborhood.

### Assigning subclass, supertype and cluster names

We first annotated each subclass with its most representative anatomical region(s) and named the subclass using the combination of its representative region(s), major neurotransmitter, and in some cases one or two marker genes. We then ordered the subclasses based on the taxonomy tree and assigned subclass IDs accordingly. Supertype names within each subclass were defined by combining the subclass name and the grouping numbers of supertypes within the subclass.

Supertype IDs were assigned sequentially based on the taxonomy tree order of subclasses and the group order of supertypes within each subclass. Cluster IDs were also assigned sequentially based on the ordering of subclasses and supertypes. And the final cluster names were assigned by combining each cluster’s ID with the name of the supertype the cluster belongs to. Based on the Allen Institute proposal for cell type nomenclature^135^, we also assigned accession numbers to cell types, as included in **Supplementary Table 7**.

### Assigning cell type identities within a modality (for cross validation) and across modalities

We performed 5-fold cross validation using different sets of marker genes: all 8,108 marker genes (**Marker gene selection section)**, 499 TF marker genes, and 20 sets of 499 randomly sampled marker genes from the 8,108-marker list. We defined the cluster centroid in each modality as the average gene expression for all the training cells within the cluster and built the Annoy KNN indices based on user specified distance metrics (cosine by default) using the chosen marker list. For the testing cells in each modality, we assigned their cell type identities by mapping them to the nearest cluster centroid using the corresponding Annoy index. This process is implemented in map_cells_knn_big function from scrattch.bigcat package, and mapping can be performed very efficiently by massive parallelization. We also used this approach for assigning cell type identities for MERFISH or any external datasets to the 10xv3 dataset as reference, using different gene lists based on the contexts. When mapping confidence was needed, we sampled 80% genes from the marker list randomly, and performed mapping 100 times. The fraction of times a cell is assigned to a given cell type is defined as the mapping probability.

### Defining transcription factor (TF) co-expression gene modules

To identify TF gene modules that are involved in regulating major cell types, we performed WGCNA analysis^136^ on 499 TF marker genes (**Supplementary Table 8**) based on their average expression at the subclass level with power = 6 and TOMType = “signed”, and detectCutHeight = 0.998. Genes in “grey” module were removed, which had poor correlation with all the other genes, and genes that were generally enriched in neurons were excluded. Genes in some modules clearly had distinct patterns and were thus further split, and they were re-ordered for better visualization.

### Defining transcription factor (TF) code along the taxonomy tree

For each node along the taxonomy tree, we computed the most discriminative TFs distinguishing all the subclasses under this node from all the subclasses under any sibling nodes, and other subclasses that also express the same combo of markers along the path from the root. If all the TF markers along the path from the root together were not specific to the given node, additional TF markers were selected to provide more specificity. We also required all selected markers at any nodes to be expressed in at least 70% of clusters within the corresponding subtree at logCPM > 2. Given such constraint, it is possible that a TF was chosen for more than one sibling nodes, in which case, we tried to select more TFs for further discrimination.

## MERFISH

### Brain dissection and freezing

Standard procedures were developed to isolate, cut, fix and pre-treat tissue to preserve macro and cellular morphology and to produce the best signal to noise ratio for MERFISH. Mice were transferred from the vivarium to the procedure room with efforts to minimize stress during transfer. If mouse body weight fell outside of the normal range (18.8 to 26.4 g), the brain was not used in the MERFISH process. Mice were anesthetized with 0.5% isoflurane. A grid-lined freezing chamber was designed to allow for standardized placement of the brain within the block in order to minimize variation in sectioning plane. Chilled OCT was placed in the chamber, and a thin layer of OCT was frozen along the bottom by brief placement of the chamber in a dry ice ethanol bath. The brain was rapidly dissected and placed into the OCT. The orientation of the brain was adjusted using a dissecting scope, and the freezing chamber containing OCT and brain were frozen in a dry ice/ethanol bath. Brains were stored at -80°C.

### Cryosectioning

The fresh frozen brain was sectioned at 10 µm on Leica 3050 S cryostats. The OCT block containing a fresh frozen brain was trimmed in the cryostat until reaching the desired starting section. Sections were collected every 200 µm to evenly cover the brain from anterior to posterior and each section was mounted onto a functionalized 20-mm coverslip treated with yellow green (YG) fluorescent microspheres (VIZGEN, #2040003)

### Fixation and dehydration

After air drying on the coverslips for 10-15 minutes, the tissue sections were loaded into a Leica Autostainer XL (Leica ST5010). They were washed in 1x PBS for 1 minute, fixed in 4% PFA for 15 minutes, washed in 1x PBS for 5 minutes 3 times, washed in 70% ethanol and then stored in 70% ethanol at 4°C. They were stored for at least one day and no more than 6 weeks before proceeding.

### Hybridization

For staining the tissue with MERFISH probes a modified version of instructions provided by the manufacturer was used. All solutions were prepared according to the instruction provided by the manufacturer. For hybridization samples were removed from the 70% ethanol and washed in a petri dish containing VIZGEN Sample Prep Buffer (VIZGEN, #20300001). Sample Prep Buffer was aspirated, and the samples were equilibrated with 5mL of VIZGEN Formamide Wash Buffer (VIZGEN, #20300002) in a humidified incubator at 37°C for 30 minutes. Formamide Wash Buffer was removed via aspiration and a 50-μl droplet of MERSCOPE Gene Panel Mix was added onto the center of the tissue section. Next, the tissue section was covered with parafilm and stored in a humidified 37°C cell culture incubator for 36-48 hours.

### Gel embedding

Parafilm covering the sections was removed and 5ml of the VIZGEN Formamide Wash Buffer was immediately added. Sections were incubated at 47°C for 30 min. Formamide Wash Buffer was aspirated and the previous step repeated. Sections were washed with VIZGEN Sample Prep Wash Buffer after the second formamide wash for 2 min. 110 µl of VIZGEN gel embedding solution (VIZGEN #20300004) with APS and TEMED was added onto the center of a Gel Slick-coated microscope slide and any excess embedding solution was gently removed.

To allow for the gel to fully polymerize the sections were incubated at room temperature for 1.5 hours. To clear the tissue the section was incubated in 5 ml of VIZGEN Clearing Solution (VIZGEN #20300003) with Proteinase K (NEB P8107S) according to the Manufacturer’s instructions for at least 24 hours or until it was clear in a humidified incubation oven at 37°C.

### Imaging

Following clearing, sections were washed twice for 5 min in Sample Prep Wash Buffer (VIZGEN, #20300001). VIZGEN DAPI and PolyT Stain (VIZGEN, 20300021) was applied to each section for 15 min followed by a 10 min wash in Formamide Wash Buffer. Formamide Wash Buffer was removed and replaced with Sample Prep Wash Buffer during MERSCOPE set up. 100 µl of RNAse Inhibitor (New England BioLabs M0314L) was added to 250 µl of Imaging Buffer Activator (VIZGEN, #203000015) and this mixture was added via the cartridge activation port to a pre-thawed and mixed MERSCOPE Imaging cartridge (VIZGEN, #1040004). 15 ml mineral oil (Millipore-Sigma m5904-6X500ML) was added to the activation port and the MERSCOPE fluidics system was primed according to VIZGEN instructions. The flow chamber was assembled with the hybridized and cleared section coverslip according to VIZGEN specifications and the imaging session was initiated after collection of a 10X mosaic DAPI image and selection of the imaging area. For specimens that passed the minimum count threshold, imaging was initiated, and processing completed according to VIZGEN proprietary protocol.

### Data analysis

Cell segmentation was performed as described previously^137^. Briefly, cells were segmented based on DAPI and PolyT staining using Cellpose^138^. Segmentation was performed on a median z-plane (4^th^ out of 7) and cell borders were propagated to z-planes above and below. The resulting cell-by-gene table was filtered to keep cells with a volume > 100 µm^3^ and < 3,000 µm^3^, that have at least 15 genes detected and contain a minimum of 40 but no more than 3,000 mRNA molecules (red dashed lines in **Extended Data Figure 2d-e**) and remove low quality cells and doublets that are outside of these ranges. Overall counts of genes were normalized by cell volume and log2 transformed. To assign cluster identity to each cell in the MERFISH dataset, we mapped the MERFISH cells to the scRNA-seq reference taxonomy. For this, the 10xv3 scRNA-seq data was subsetted to only genes common to both datasets. Our mapping method (as described in **Assigning cell type identities** section) finds the nearest cluster centroid in the scRNA-seq reference dataset for a query data point with the correlation of shared genes as distance metric. The cluster label of the nearest neighbor was assigned as mapped label.

Bootstrapping was conducted with 80% subsampling of marker genes to make label assignment robust.

### CCF registration

To facilitate alignment of MERFISH sections to the CCF, we assigned each cell from the scRNA-seq dataset to one of these major regions: CB, CTXsp, HB, HPF, HY, isocortex, LSX, MB, OLF, PAL, sAMY, STRd, STRv, TH and HB. This delineation was driven by the level of region-specific dissection for the scRNA-seq experiments as well as the cell type specificity of regions. Because of the more gradient transition of cell type composition between cortical regions, the specificity of cortical plate regions is limited to isocortex, OLF and HPF despite more granular dissection regions. Each cluster in the scRNA-seq dataset was assigned to the region the majority of cells were derived from. We identified anchor clusters we used for region annotation of the MERFISH data. These clusters were defined as a) having more that 30% of all cells in one region and b) more than 20 cells in a MERFISH section. In addition to that we used ependymal and choroid plexus cells to label the ventricles and identified specific clusters of oligodendrocytes that were enriched in white matter tracts. To account for clusters that were found at low frequency in regions outside its main region we calculated for each cell its 50 nearest neighbors in physical space and reassigned each cell to the region annotation dominating its neighborhood. Next, we used that same approach to assign each cell mapped to a non-anchor cluster to the region annotation dominating its immediate surrounding. The resultant label maps were used as input to our registration tool to find for each section its approximate location along the anterior to posterior axis of the brain as well as any offsets in pitch and yaw introduced during sectioning.

Registration was performed at 10-µm in-plane resolution. For each section, an anatomical reference image was created by aggregating the number of detected spots within a 10x10 µm grid for each gene probe. A single image was created across all probes by taking the maximum count for each grid unit. The midline was manually determined by annotating the most dorsal and most ventral point. These points were then used to compute a rigid transform to rotate the section upright and center in the middle. This set of rectified images were stacked in sequencial order to create an initial configuration for registration.

Alignment to the Allen CCFv3 was performed by matching the above-mentioned scRNA-seq derived region labels to their corresponding anatomical parcellation of the CCF. A label map was generated for each region by aggregating the cells assigned to that region within a 10x10 µm grid, transformed to the initial configuration using the computed rigid transforms. Using the corresponding anatomical labels, the ANTS registration framework was used to establish a 2.5D deformable spatial mapping between the MERFISH data and the CCF via three major steps: 1) A 3D global affine (12 dof) mapping was performed to align the CCF into the MERFISH space.

This generated resampled sections from the CCF that provided section-wise 2D target space for each of the MERFISH sections. Since the CCF is a continuous label set with isotropic voxels, this avoids interpolation artifacts that can result if resampling is performed on the MERFISH data instead, which has large section gaps, and can contain missing sections. 2) After establishing a resampled CCF section for each MERFISH section, 2D affine registrations were performed to align each MERFISH section to match the global anatomy of the CCF brain. This addressed misalignments from the initial manual stacking of the MERFISH sections using the midline and provided a global mapping to initialize the local deformable mappings. 3) Finally, a 2D multi-scale, symmetric diffeomorphic registration (step size = 0.2, sigma = 3) was used on each section to map local anatomic differences between the corresponding MERFISH and CCF structures in each section. Global and section-wise mappings from each of these registration steps were preserved and concatenated (with appropriate inversions) to allow point-to-point mapping between the original MERFISH coordinate space and the CCF space.

## ACKNOWLEDGEMENTS

We are grateful to the Transgenic Colony Management, Lab Animal Services, Molecular Biology, Spatial Transcriptomics, and Histology teams at the Allen Institute for technical support. The research was funded by the U19MH114830 grant from National Institute of Mental Health to H.Z. and X.Z., under the BRAIN Initiative of National Institutes of Health (NIH). The content is solely the responsibility of the authors and does not necessarily represent the official views of NIH and its subsidiary institutes. X.Z. is a Howard Hughes Medical Institute investigator. This work was also supported by the Allen Institute for Brain Science. The authors thank the Allen Institute founder, Paul G. Allen, for his vision, encouragement, and support.

## Author Contributions

Conceptualization: H.Z. Data analysis lead and coordination: Z.Y. Data generation (scRNA-seq): Z.Y., C.T.J.vV., D.M., A.A., E.B., D.B., D.C., T.C., M. Clark, K.C., L. Ellingwood, A.G., J. Gloe, J. Gray, N.G., J. Guzman, D.H., W.H., M. Kroll, K.L., R.M., E.M., T.N.N., T.P., C.R., N.S., J.S., M.T., A.T., H.T., K. Wadhwani, K. Ward, B. Levi, N.D., K.A.S., B.T., H.Z. Data processing and analysis (scRNA-seq): Z.Y., C.T.J.vV., C.L., J. Goldy, A.B.C., R.C., S.C., T.D., J. Gould, K.A.S., B.T., H.Z. Data generation (MERFISH): M. Kunst, M.Z., D.M., W.J., J. Campos, N.M., A.R., N.V.C., J.W., X.Z., H.Z. Data processing and analysis (MERFISH): Z.Y., M. Kunst, M.Z., D.M., W.J., M. Chen, J. Close, S.D., J. Gee, A.L., B. Long, Z.M., C.S., J.W., L.N., X.Z., H.Z. Project management: P.B., C.L.T., C.P., L.K., S.M.S., K.A.S. Management and supervision: Z.Y., C.T.J.vV., D.M., J. Close, J. Gee, B. Long, B. Levi, C.F., C.L.T., S.M., N.D., S.M.S., L. Esposito, M.J.H., J.W., L.N., K.A.S., B.T., X.Z., H.Z. Manuscript writing and figure generation: Z.Y., C.T.J.vV., M. Kunst, H.Z. Manuscript review and editing: Z.Y., C.T.J.vV., M. Kunst, K.J., M.J.H., L.N., B.T., X.Z., H.Z.

## Additional Information

Correspondence and inquiries about data and materials should be addressed to: H.Z. or Z.Y.

## Data Availability

The data were generated under the BRAIN Initiative Cell Census Network (BICCN, www.biccn.org, RRID:SCR_015820) and are accessible through Neuroscience Multi-omic Data Archive (NeMO, https://nemoarchive.org/, with identifiers nemo:dat-kjseyhn and nemo:dat- 5ue6x8t) and Brain Image Library (BIL, https://www.brainimagelibrary.org/index.html). The AIBS 10x scRNA-seq dataset reported in this study will be available at NeMO site https://data.nemoarchive.org/publication_release/ under bundle name ‘Zeng_transcriptome_Allen_10x_cells_wholebrain_2023’. The AIBS MERFISH dataset will be available at BIL under DOI https://doi.org/10.35077/g.610.

## Code Availability

Data analysis code used in the manuscript, R package scrattch.bigcat, is available via github https://github.com/AllenInstitute/scrattch.bigcat.

---

**Extended Data Figure 1.**
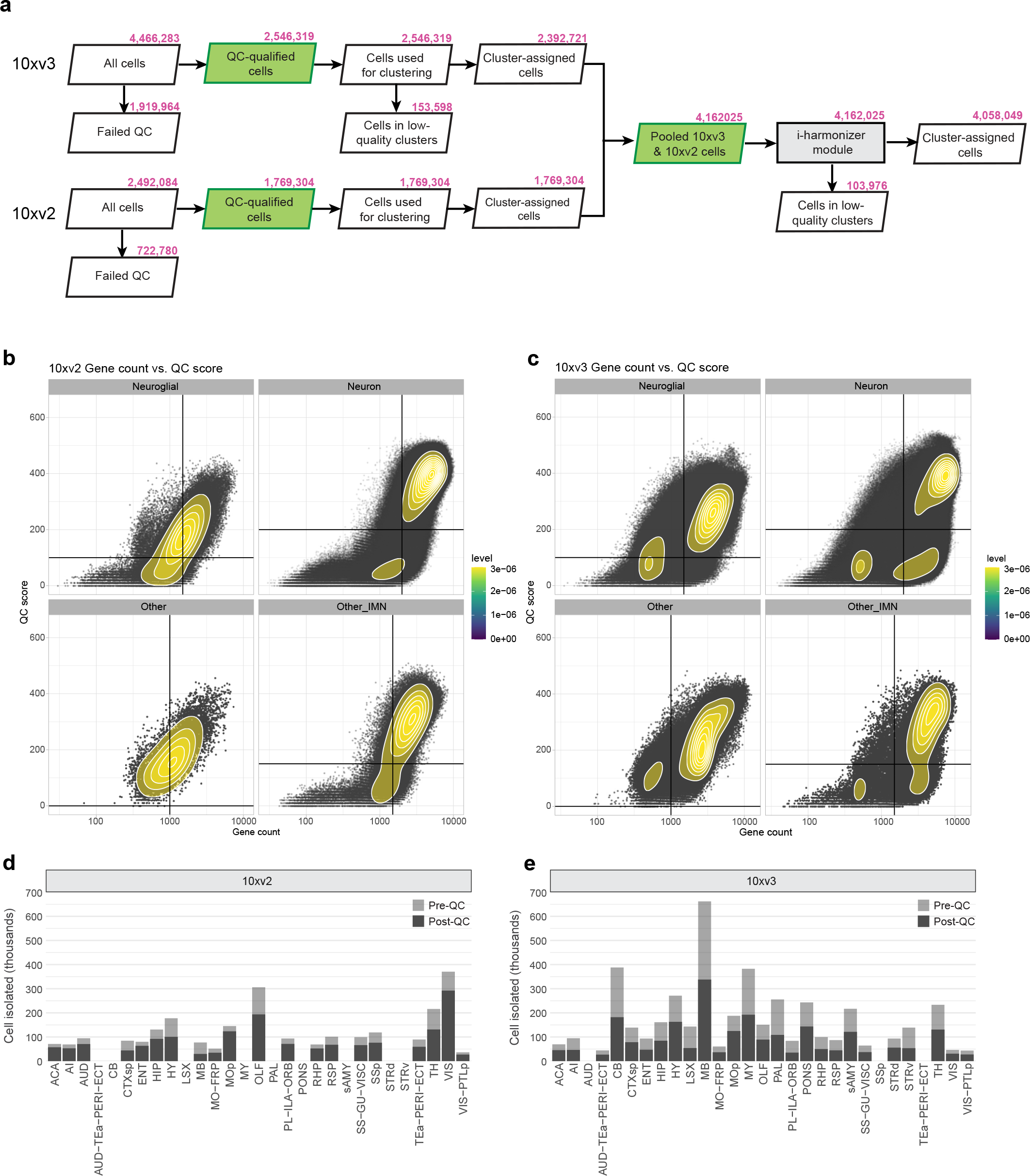
scRNA-seq data analysis workflow. (a) Number of cells at each step in the scRNA-seq data analysis pipeline. The identification of doublets and low-quality clusters is described in more detail in Methods. The 10xv2 and 10xv3 data were first QC-ed and analyzed separately. After initial clustering the datasets were combined and QC-ed again before and after joint clustering. **(b-c)** Gene count and qc score thresholds used for each of the four major cell populations (neuroglial cells, neurons, immature neurons and granule cells, and other) on the 10xv2 (b) and 10xv3 (c) datasets. **(d-e)** Number of cells isolated from dissection ROI’s (pre-QC) and number of cells passing QC (post-QC) for 10xv2 (d) and 10xv3 (e) datasets. We didn’t profile LSX, STR, sAMY, PAL, Pons, MY, and CB by 10xv2. Some regions were collected using different dissections between 10xv2 and 10xv3, but all regions were covered by 10xv3.

**Extended Data Figure 2.**
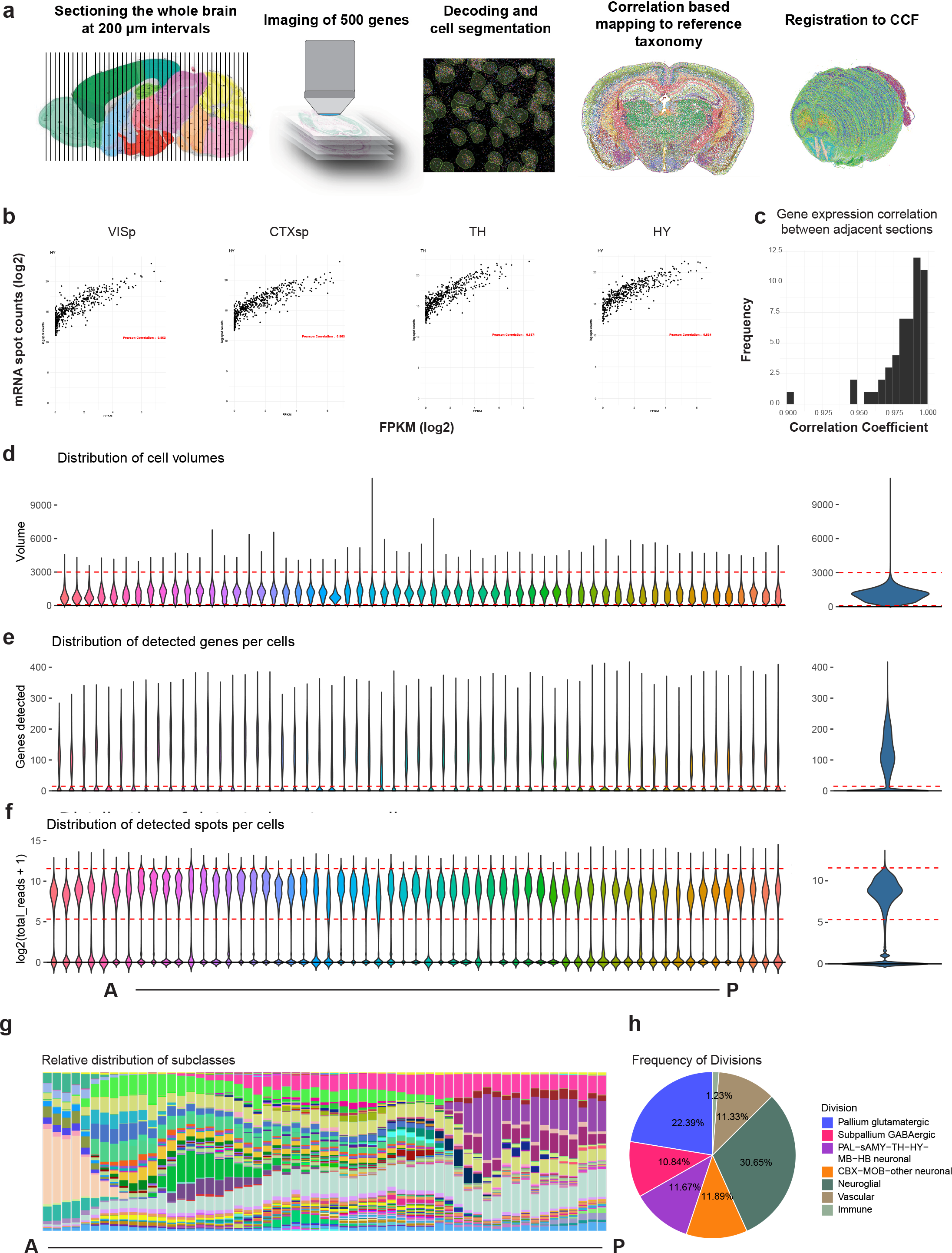
MERFISH data generation, data processing and summary of results. (a) Workflow for generating and processing MERFISH data. **(b)** Correlation of gene detection between MERFISH and bulk RNA-sequencing for four different brain regions. **(c)** Histogram displaying the distribution of gene detection correlation between adjacent MERFISH sections. **(d-f)** Violin plots displaying distribution of cell volumes (d), gene detection (e), and mRNA molecule detection (f) for individual sections ordered from anterior to posterior (left panel) or cumulative distribution for the whole brain (right panel). Red dashed lines indicate cutoff for filtering. **(g)** Cumulative histogram showing the relative contribution of each subclass to each section ordered from anterior to posterior. **(h)** Pie chart showing the proportion of cells in each major division across the whole brain.

**Extended Data Figure 3.**
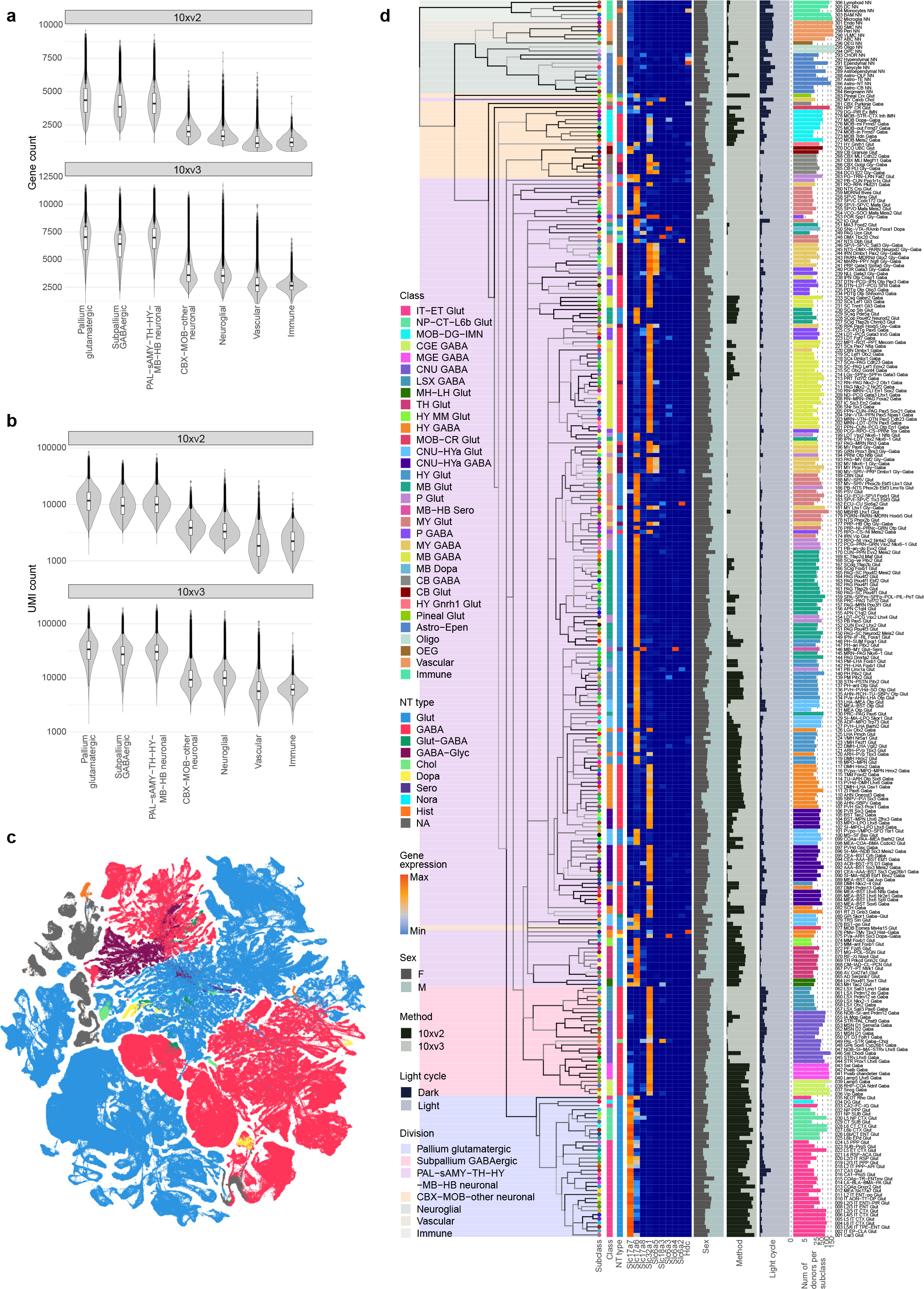
Transcriptomic cell type taxonomy of the whole mouse brain with additional metadata information. (a-b) Number of genes (a) or number of UMI’s (b) detected per cell in 10xv2 (top) or 10xv3 (bottom) datasets for each major cell division. The data shown is post-QC. **(c)** UMAP representation of all cell types colored by neurotransmitter (NT) type. NT type color code is the same as shown in (d). **(d)** The transcriptomic taxonomy tree of 306 subclasses organized in a dendrogram (same as Figure 1a). The color blocks divide the dendrogram into major cell divisions. From left to right, the bar plots represent cell class assignment, NT type assignment, heatmap showing expression of major neurotransmitter marker genes, sex distribution, platform distribution, light-dark distribution of profiled cells, and number of donors that contributed to each subclass.

**Extended Data Figure 4.**
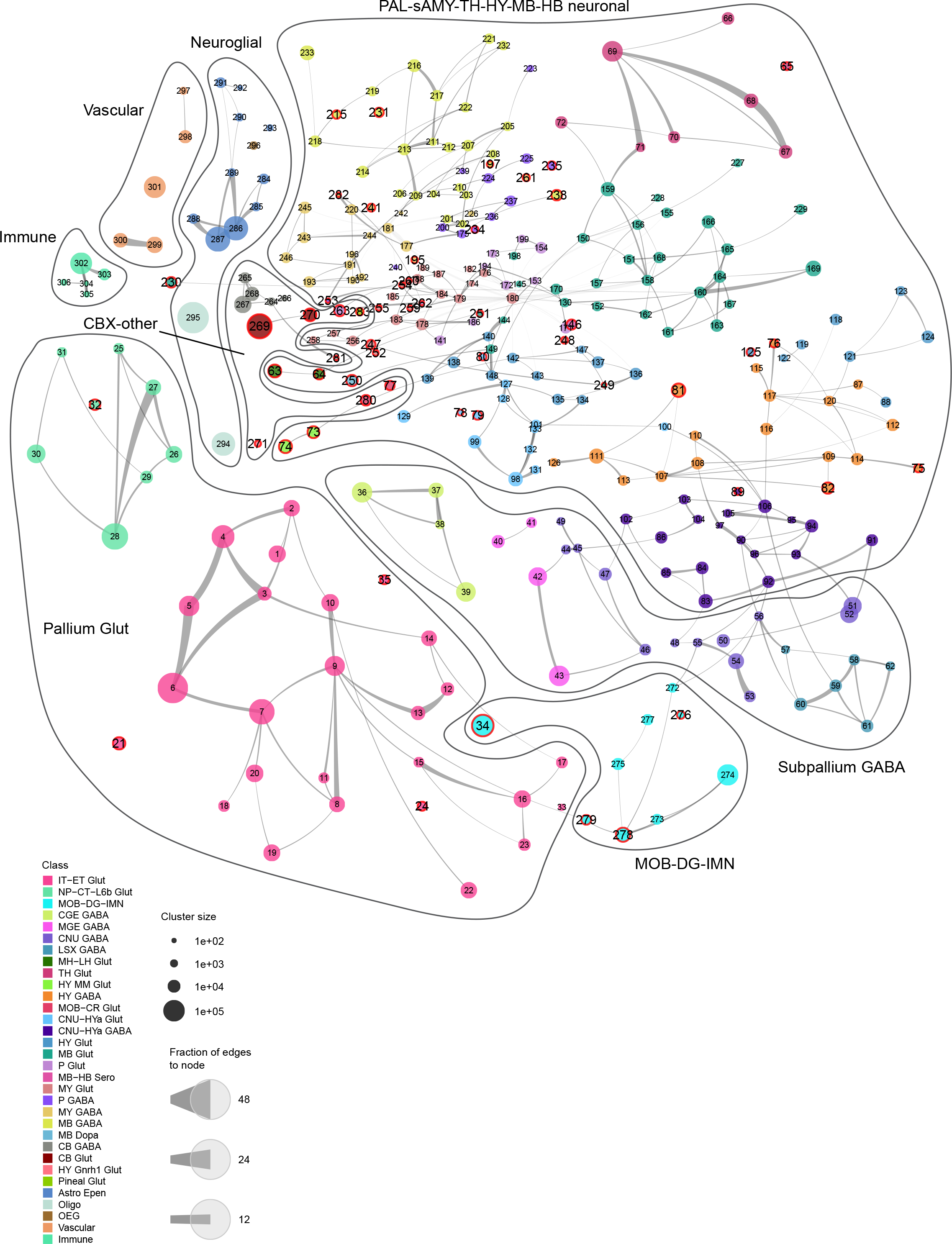
Constellation plot of the global relatedness between subclasses. Each subclass is represented by a disk, labeled by the subclass ID and positioned at the subclass centroid in UMAP coordinates shown in Figure 1d. The size of the disk corresponds to the number of cells within each subclass, and the edge weights correspond to the fraction of shared neighbors (see Methods) between subclasses. Each subclass is colored by the class it belongs to. Curved line bubbles drawn around subclasses outline the major divisions. Distinct subclasses are highlighted by the red rings around the disks.

**Extended Data Figure 5.**
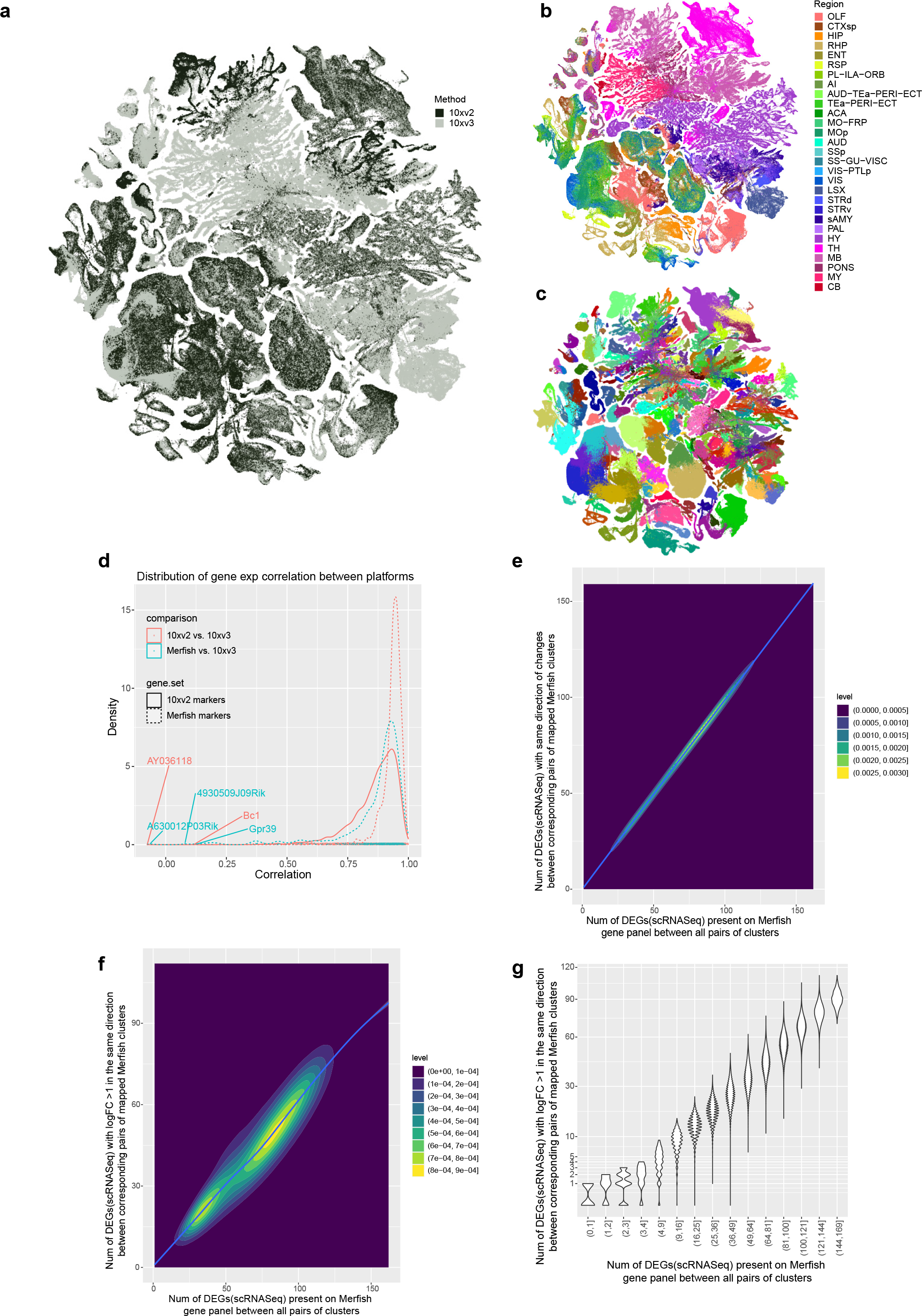
Validation of data integration across 10xv2, 10xv3, and MERFISH datasets. (a-c) UMAP representation of all cell types colored by profiling platform (a), region (b), and subclass (c). Other than the regions only profiled by 10xv3 (LSX, STR, sAMY, PAL, Pons, MY), the cells from both platforms integrate very well. Cell types in isocortex and HPF have a lot more 10xv2 cells, consistent with our sampling plan. **(d)** Correlation of gene expression between 10xv2 and 10xv3 and between 10xv3 and MERFISH. For each gene, we computed the Pearson correlation of its average expression in each cluster across clusters between 10xv2 and 10xv3, and the correlation between 10xv3 and MERFISH. For 10xv3 and MERFISH comparison, distribution of the correlation values of all 500 genes in the MERFISH panel is shown. For 10xv3 and 10xv2 comparison, we show the correlation of 5383 marker genes based on 10xv2, and 466 10xv2 marker genes that are also present on the MERFISH gene panel (the 34 MERFISH genes not shown are expressed in clusters not profiled by 10xv2). **(e)** 2D density plot showing on the X-axis the number of DEGs (based on 10xv3 dataset) present on the MERFISH gene panel between all pairs of clusters, and on the Y-axis the number of such DEGs showing the same direction of changes between corresponding pairs of mapped MERFISH clusters. Almost all the DEGs between all pairs of clusters show the same direction of changes between 10xv3 and MERFISH. **(f)** 2D density plot showing on the X-axis the number of DEGs (based on 10xv3 dataset) present on the MERFISH gene panel between all pairs of clusters, and on the Y-axis the number of such DEGs showing the same direction of changes, and logFC > 1 between corresponding pairs of mapped MERFISH clusters. About 60% of DEGs between all pairs of clusters based on 10xv3 show significant fold change (FC) in MERFISH. **(g)** Similar analysis as in (f) but shown as violin plot by binning the number of 10xv3 DEGs present on the MERFISH gene panel on the X-axis, with better resolution on closely related pairs with four or fewer DEGs present on MERFISH gene panels.

**Extended Data Figure 6.**
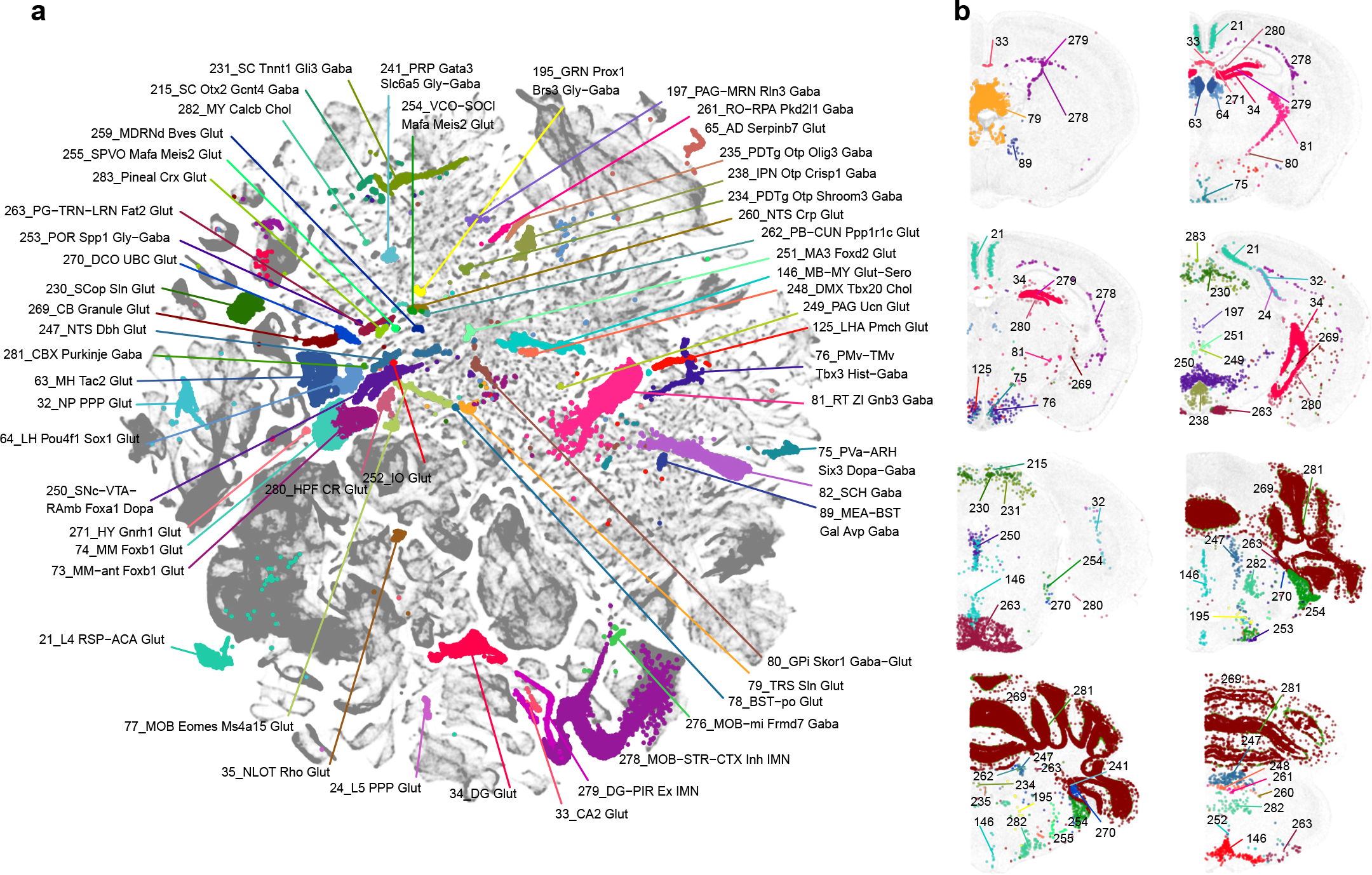
Highly distinct neuronal types across the brain. UMAP representation **(a)** and representative MERFISH sections **(b)** of highly distinct subclasses across the brain, colored by subclass.

**Extended Data Figure 7.**
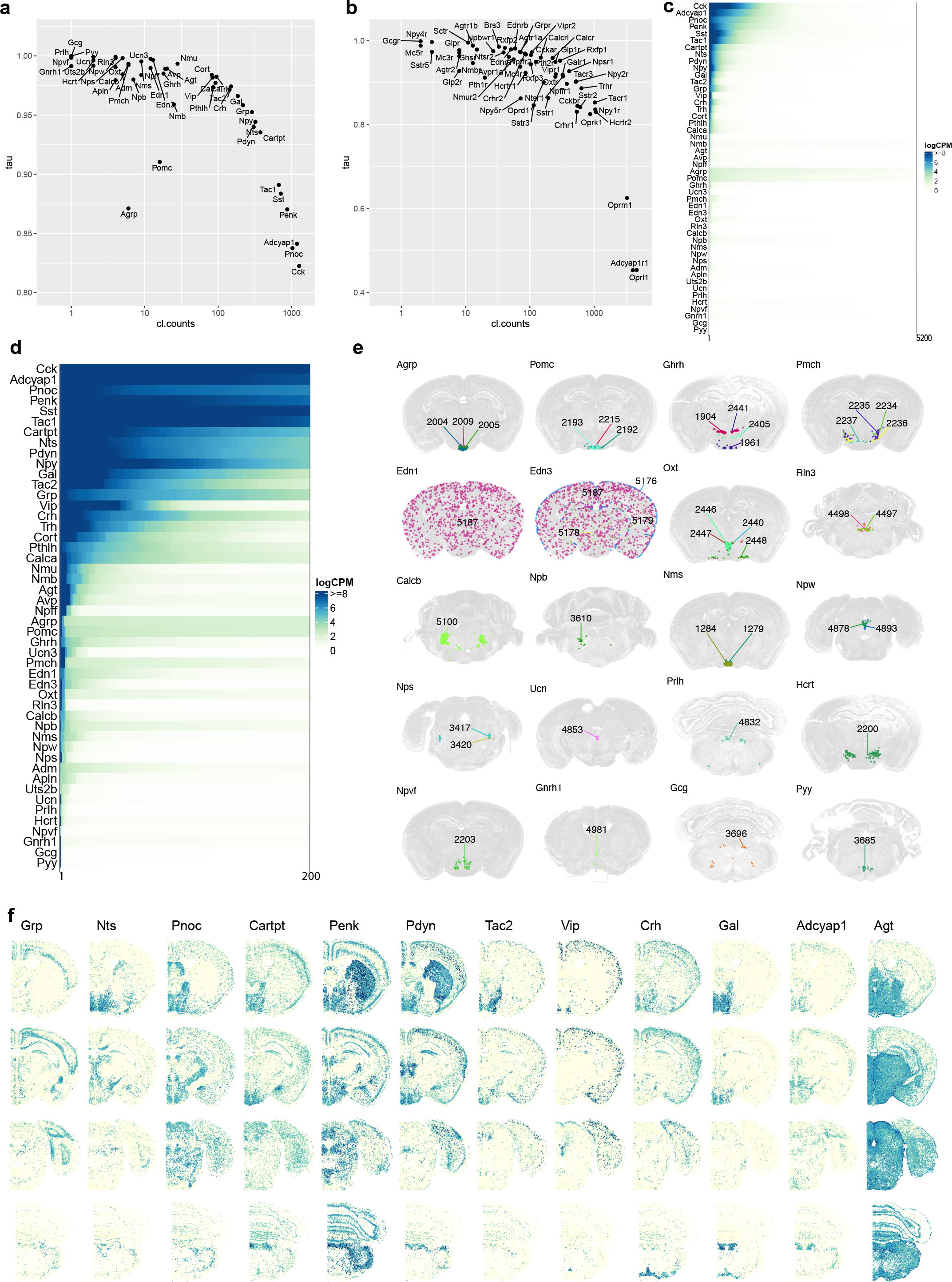
Neuropeptide distribution across the whole mouse brain. (a) Scatter plot of Tau score over the number of clusters each neuropeptide is expressed in at the level of logCPM > 3. The Tau score is a measurement of cell type specificity, which varies from 0 to 1 where 0 means uniformly expressed and 1 means highly specific to one type. **(b)** Scatter plot of Tau score over the number of clusters each peptide-liganded G-protein coupled receptor (GPCR) gene is expressed in at the level of logCPM > 3. **(c)** Expression level of neuropeptide (logCPM) per cluster. For each neuropeptide along the Y axis, clusters are sorted from the highest to lowest mean gene expression level along the X axis. **(d)** Expression level of neuropeptide (logCPM) per cluster. For each neuropeptide along the Y axis, clusters are sorted from the highest to lowest mean gene expression level along the X axis. For each gene, only the top 200 highest-expressing clusters out of 5,200 clusters are shown. **(e)** Representative MERFISH sections highlighting the spatial location of clusters expressing each of the 20 highly cell-type-specific neuropeptide genes (expressed in 8 or fewer clusters). **(f)** Representative MERFISH sections showing the expression of the neuropeptides present on the MERFISH gene panel that are widely expressed.

**Extended Data Figure 8.**
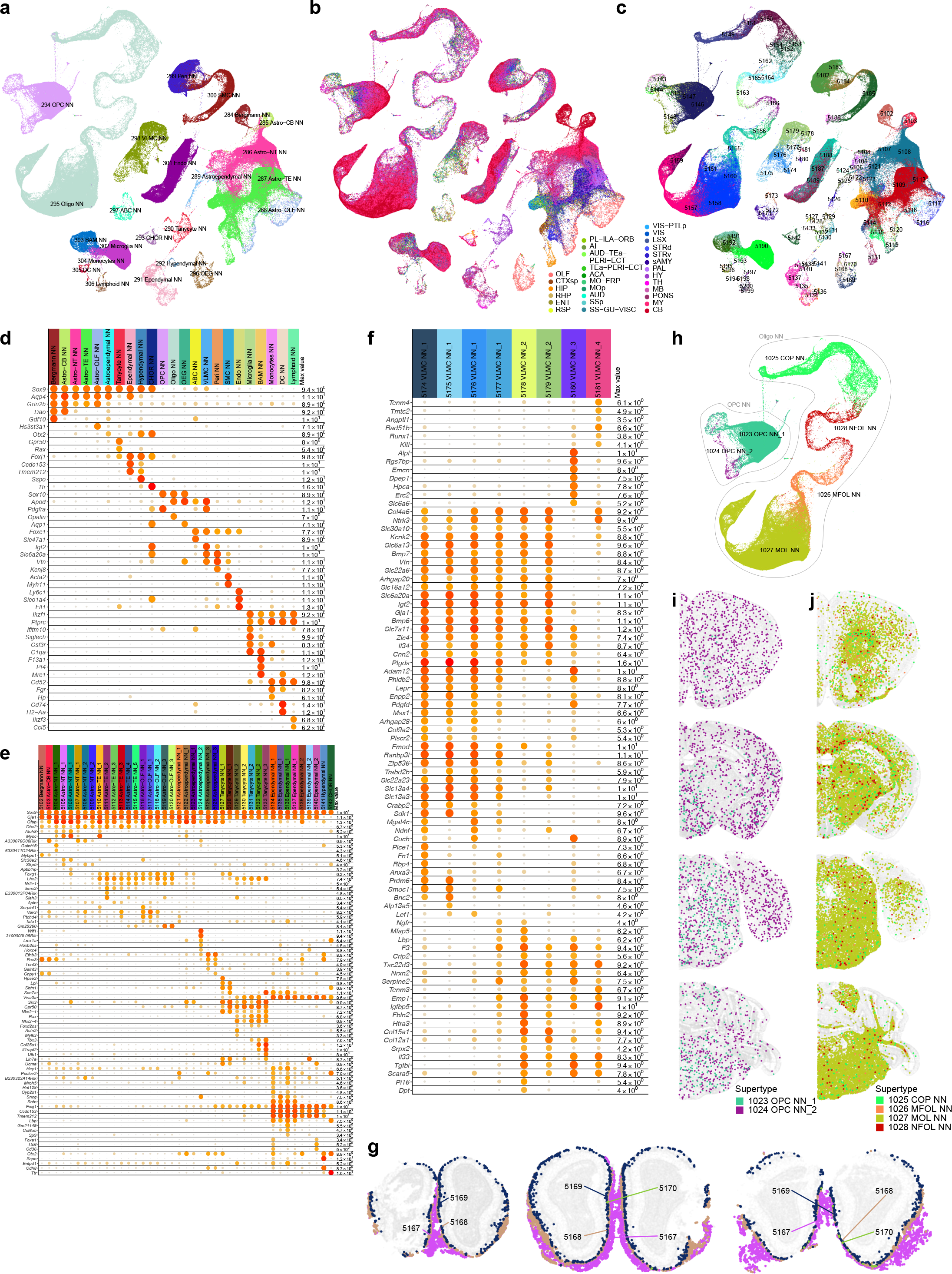
Additional non-neuronal UMAPs and marker genes. (a-c) UMAP representation of non-neuronal cell types colored by subclass (a), region (b), and cluster (c). **(d)** Dot plot showing marker gene expression in non-neuronal subclasses. Dot size and color indicate proportion of expressing cells and average expression level in each subclass, respectively. **(e)** Dot plot showing marker gene expression in all clusters in the Astro-Epen class. Dot size and color indicate proportion of expressing cells and average expression level in each cluster, respectively. **(f)** Dot plot showing the marker gene expression in VLMC clusters. Dot size and color indicate proportion of expressing cells and average expression level in each cluster, respectively. **(g)** Representative MERFISH sections showing the spatial gradient of OEG clusters. **(h)** UMAP representation of OPCs and oligodendrocytes colored and labeled by supertype. **(i-j)** Representative MERFISH sections showing the spatial distribution of OPC (i) and Oligo (j) supertypes.

**Extended Data Figure 9.**
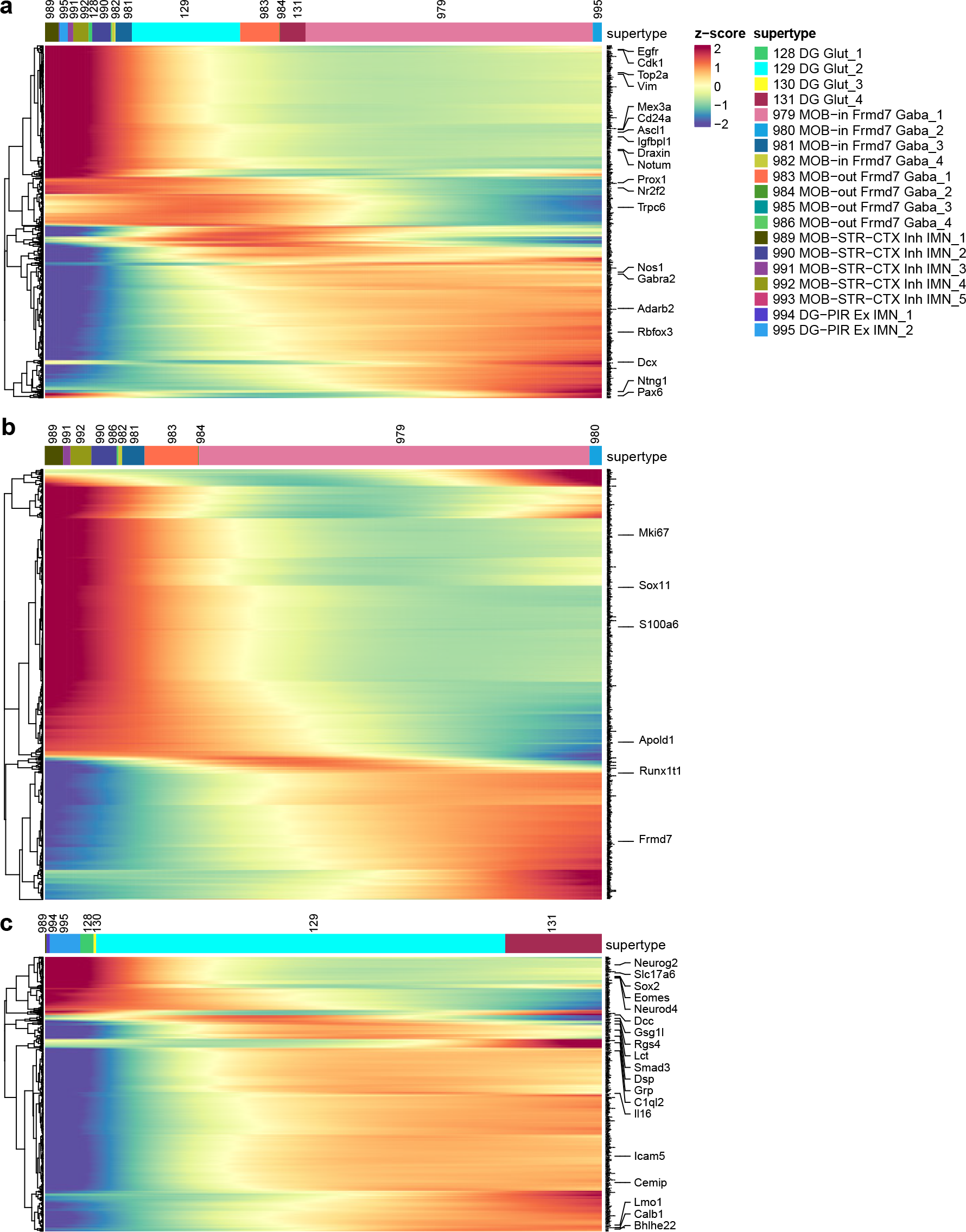
Gene expression patterns in immature neuron populations. (a) Heatmap showing the gene expression changes as immature neurons transition to mature cell types, conserved between DG and MOB cell type development. Key markers at each stage of development are highlighted. **(b)** Heatmap showing the gene expression changes as immature neurons transition to mature cell types, specific to MOB cell types. **(c)** Heatmap showing the gene expression changes as immature neurons transition to mature cell types, specific to DG cell types.

**Extended Data Figure 10.**
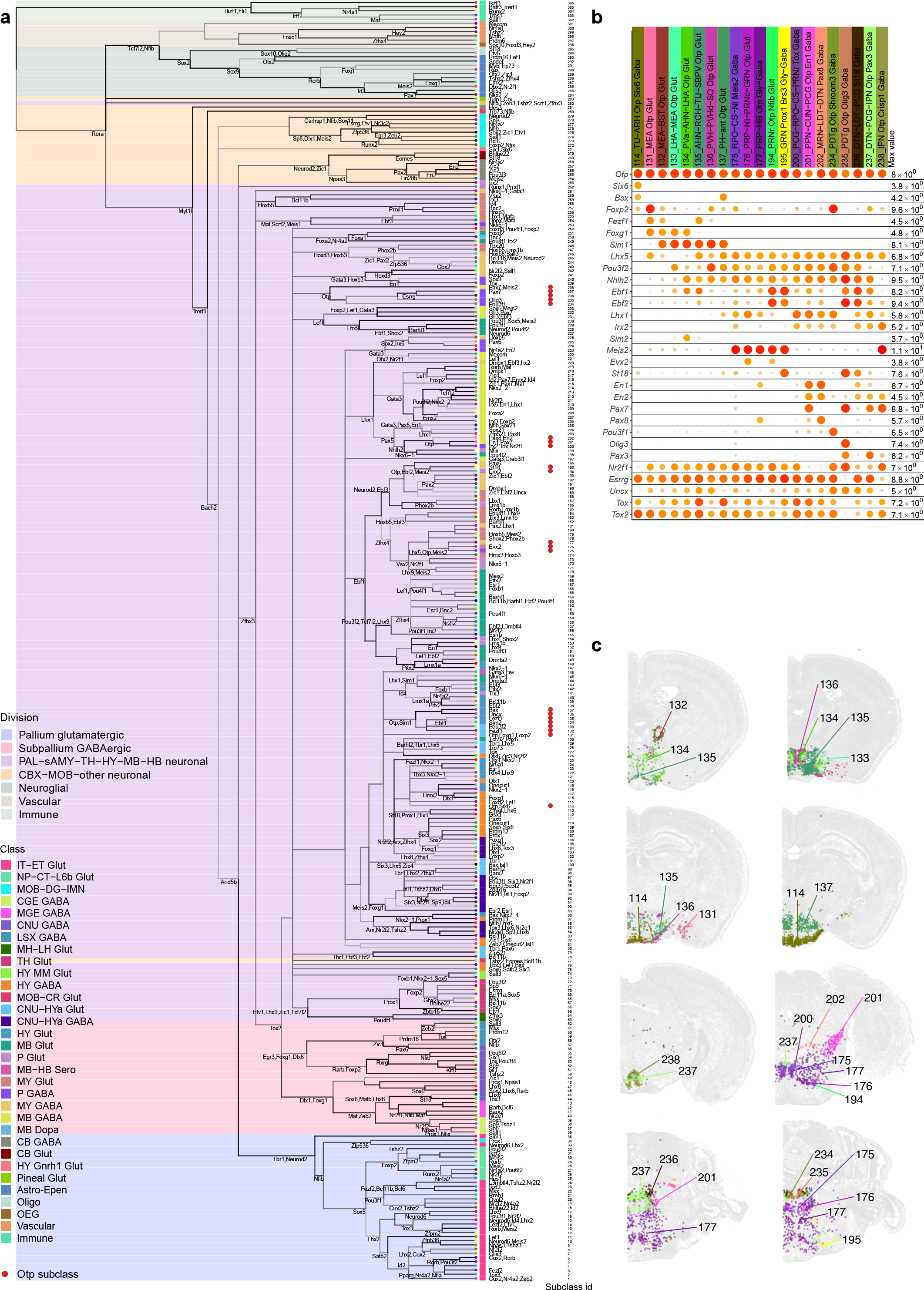
Transcription factor code. (a) The transcriptomic taxonomy tree of 306 subclasses organized in a dendrogram (same as Figure 1a). The color blocks divide the dendrogram into major cell divisions. The color bars denote classes. Key transcription factors are annotated for nodes and subclasses on the tree. Red dots mark the Otp expressing subclasses described in panels (b) and (c). **(b)** Gene expression dot plot of Otp expressing subclasses. Dot size and color indicate proportion of expressing cells and average expression level in each subclass, respectively. **(c)** Representative MERFISH sections highlighting the Otp expressing subclasses.

**Extended Data Figure 11.**
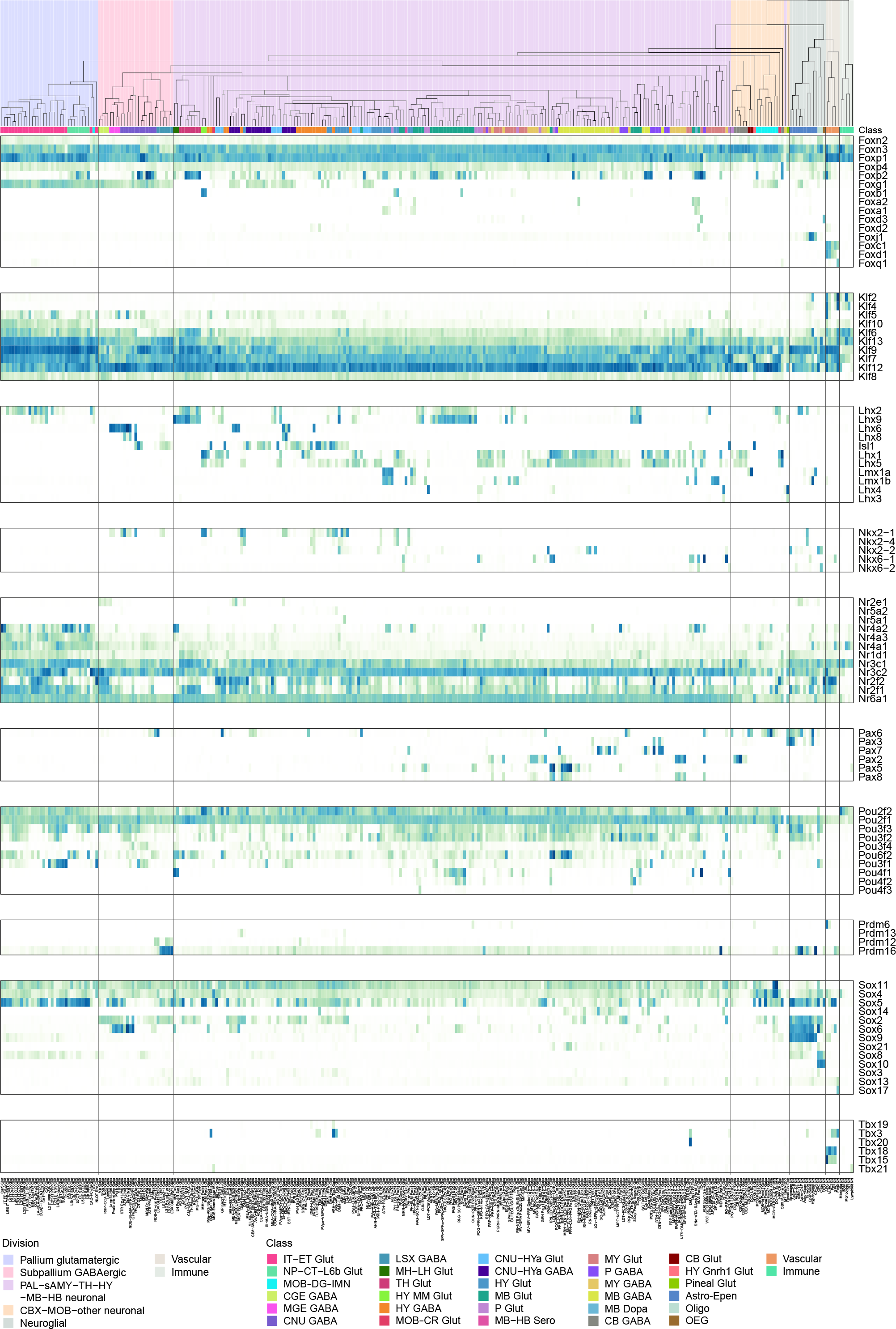
Transcription factor families. Expression of key TFs for each subclass in the taxonomy tree, organized by TF gene families. The color blocks divide the dendrogram into major cell divisions. The color bars denote classes.

**Extended Data Figure 12.**
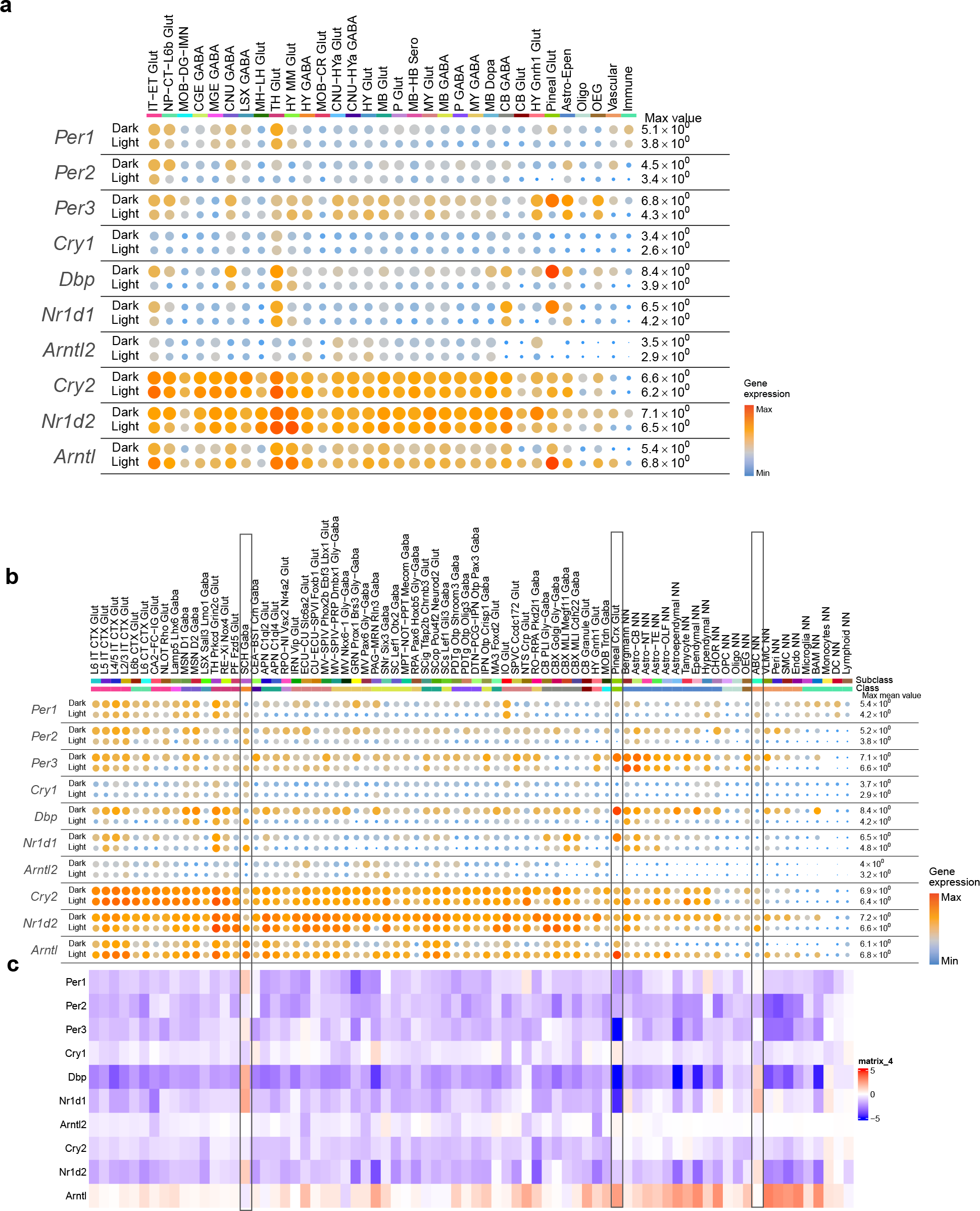
Circadian cycle associated expression changes in clock genes. (a-b) Dot plot showing the expression of clock genes in light-phase and dark-phase cells within each cell class (a) or selected subclasses that have any clock genes with fold change logFC > 1 between light and dark phases (b). Dot size and color indicate proportion of expressing cells and average expression level in each class or subclass, respectively. **(c)** Heatmap showing the logFC difference between light and dark phases for clock genes in selected subclasses as in (b).

**Supplementary Table 1. Allen Mouse Brain Common Coordinate Framework version 3 (CCFv3) regional ontology.** Adopted from Wang et al, 2020.

**Supplementary Table 2. RNA-seq specimen information.** All donors used in this study are listed, with associated metadata including sex, age, genotype, light/dark cycle phase, etc. From one donor multiple regions could be dissected (“roi.1”, “roi.2”, “roi.3”) or multiple FACS gating plans (“facs_population_plan.1-3”) were used.

**Supplementary Table 3. RNA-seq cell sampling per region.** Number of cells sampled for each dissected region using 10xv2 or 10xv3 platform. ROI (region of interest) is the brain region combination for the 10x profiling.

**Supplementary Table 4. RNA-seq quality control thresholds used for each cell class.** The first tab has the gene count and qc score thresholds for each cell class, the second tab has the list of genes used to calculate the qc score.

**Supplementary Table 5. Marker gene list.** The list of 8,108 differentially expressed genes (DEGs) combined from the top 15 differentially expressed genes in both directions between all pairs of clusters, which was used for imputation, PCA dimensionality reduction and 2D/3D UMAP computation.

Supplementary Table 6. MERFISH 500-gene panel used in Vizgen MERSCOPE platform.

**Supplementary Table 7. Cell type annotation.** Detailed information for each cluster, including membership in broader categories (supertype, subclass, class, division and neighborhood), NT type, NT type combo, major NT marker genes, major neuropeptides, main dissection region, tentative anatomical annotation, number of 10xv2 and 10xv3 cells, relative proportions between sexes and light/dark conditions, accession numbers to cell types, and marker genes. Note that the tentative anatomical annotations are tentative and incomplete, and they will need to be refined in the future.

**Supplementary Table 8. Transcription factor marker gene list.** The first tab shows the 499 TF marker genes contained within the 8,108 DEG list. The second tab shows the TF gene modules shown in Figure 5d.

